# RNA editing fine-tunes transcriptional plasticity and enhances the adaptability of *Myzus persicae* to diverse host plants

**DOI:** 10.1101/2025.11.30.691374

**Authors:** Wenyuan Yu, Jun Wu, Rong Hu, Zhimou Lin, Peiyu Yang, Zhichao Hu, Pengshuai Peng, Shuai Zhan, Jean-Christophe Simon, Saskia A. Hogenhout, Gang Wu, Shuangxia Jin, Yazhou Chen

**Affiliations:** Hubei Hongshan Laboratory, Wuhan, 430070, China; Hubei Insect Resources Utilization and Sustainable Pest Management Key Laboratory, College of Plant Science and Technology, Huazhong Agricultural University, Wuhan, 430070, China; Key Laboratory of Plant Design, Center for Excellence in Molecular Plant Sciences, Chinese Academy of Sciences, Shanghai, China; INRAE (National Institute of Agriculture, Food and Environment), UMR IGEPP, Le Rheu, France; Crop Genetics, John Innes Centre, Norwich Research Park, Norwich, NR4 7UH, UK; National Key Laboratory of Crop Genetic Improvement, Huazhong Agricultural University, Wuhan, Hubei 430070, P. R. China

**Keywords:** herbivorous insects, RNA editing, host adaptation, transcriptional plasticity, generalism

## Abstract

Parthenogenetic organisms can adapt to diverse and rapidly changing environments despite limited standing genetic variation, but the molecular mechanisms enabling such adaptability remain unclear. Here, we investigate this question in the green peach aphid (*Myzus persicae*, GPA), an asexual generalist herbivore capable of colonizing a wide range of host plants without underlying genetic change. By integrating RNA-seq and DNA-seq from genetically identical clones reared on different hosts, we identified 1,368 high-confidence RNA editing sites (RES), many of which exhibited strong host-dependent differences in editing levels. These RES encompassed all 12 possible nucleotide substitutions, with A-to-I, C-to-U, and U-to-C conversions being particularly prevalent. Editing at several sites shifted dynamically within 48 hours of host transfer and often coincided with changes in transcript abundance. Functional knockdown of the editing genes *ADAR1* and *ADAR2* reduced editing at specific sites, diminished fecundity, and abolished aphid survival on new hosts. Comparative analyses across aphid species reveal that although the biochemical signatures of A-to-I editing are deeply conserved, the genomic locations and functional roles of edited transcripts are species specific, reflecting lineage-specific adaptive strategies. Together, our findings demonstrate that RNA editing provides a rapid, reversible mechanism for environmental acclimation in GPA, enabling adaptive phenotypic plasticity even in the absence of genetic variation. This work establishes RNA editing as a key molecular basis for the remarkable ecological generalism of clonal organisms.

**Significance statement:** A major question in evolutionary biology is how clonal organisms with little genetic variation rapidly adapt to diverse, changing environments. Here we show that RNA editing provides a mechanism enabling the parthenogenetic green peach aphid (*Myzus persicae*) to colonize a wide range of host plants. RNA editing generates transcriptomic diversity, responds quickly to host shifts, and is essential for adaptation, as silencing ADAR genes disrupts A-to-I editing, reduces fitness, and prevents survival on new hosts. Comparative analyses across aphid species reveal that although the biochemical features of A-to-I editing are conserved, its genomic deployment and functional outcomes differ among species, reflecting lineage-specific strategies. These findings demonstrate that RNA editing creates a dynamic regulatory layer that offsets constraints of asexual reproduction.

## Introduction

Standing genetic variation fuels adaptation, as preexisting mutations that change phenotypes can be selected for and spread through the population when advantageous (1). However, whether genetic mutations alone are sufficient for an organism to adapt to rapidly changing environments remains unclear. In particular, this concept is challenged by parthenogenetic or clonal organisms that are unlikely to accumulate favorable mutations in a short period but can plastically produce alternative phenotypes in response to environmental change.

Aphids are small sap-feeding insects of the order Hemiptera and are highly specialized plant feeders that often transmit plant pathogens, especially viruses (2). Aphid life cycles typically encompass alternation of sexual and parthenogenetic stages. Sexual reproduction promotes recombination and gene shuffling, ultimately enriching genetic variation (3). However, many aphid lineages have lost the sexual phase in their life cycle and reproduce parthenogenically (4). Under conditions such as the absence of winter hosts on which sexual reproduction occurs and/or living in warm regions, parthenogenesis becomes the sole reproductive strategy year-round. Aphids produced parthenogenetically are genetically identical across the vast majority of the genomes (4, 5), and consequently, genetic variation in asexual populations is extremely low (6). Nevertheless, asexual aphids can rapidly adapt to harsh environments in both laboratory and field settings (7–10), for example, by acquiring resistance to insecticides (11) or adapting to resistant host plants (12). The mechanisms underlying such adaptability, likely independent of preexisting genetic variation, remain largely unknown.

In addition to genetic variations, nucleotide variations in RNA molecules generated by RNA editing have increasingly been recognized to promote insect adaptation (13, 14). RNA editing in insects contributes to caste determination (15), task performance (16), wing dimorphism (17), temperature adjustment (18, 19), and insecticide resistance (20–23). RNA editing modifies nucleotides in RNA molecules, leading to a significant diversity of transcripts in an environment-responsive manner. The conversion of one type of nucleotide to any of the other three types theoretically produces 12 different editing types in primary RNA transcripts (24). Consequently, editing results in amino acid changes in the encoded proteins (25), altered secondary structure of RNAs (26), replaced splice consensus sites (22, 27), and modified RNA fate (28). The most common editing event in insects is the deamination of adenosine into inosine (A>I) by Adenosine Deaminase Acting on RNA (ADAR) (26, 29). Inosines (Is) are recognized as guanosines (Gs) by the translational machinery and the reverse transcriptase used in protocols of RT-PCR and RNA-sequencing, resulting in A-to-G (A>G) substitution (30). C-to-U (C>U) editing mediated by cytidine deaminase Apolipoprotein B mRNA Editing Enzyme, Catalytic Polypeptide 1 (APOBEC-1), U-to-C (U>C) (31, 32), and G-to-A (G>A) by *trans*-amination (33) are less frequently observed in insects so far examined (24).

To investigate the RNA editing in aphids, we focused on the green peach aphid (GPA), *Myzus persicae*, a highly polyphagous generalist capable of colonizing a broad range of host plants and transmitting more than 100 plant viruses (34). GPA has gained resistance to most classes of insecticide, making it one of the most widely and strongly resistant insect species worldwide (11, 35, 36). GPA has both sexual and parthenogenetic stages in the life cycle. Sexual reproduction occurs in autumn on peach trees, and the parthenogenetic generations feed on a wide number of secondary host species, including many economically important crops. In many areas where peach trees are absent or the climate is warm, the life cycle is simplified to continual parthenogenesis throughout the year, leading to rapidly increasing population and escalating field damage. Unlike some herbivorous insects that are generalists at the species level but show plant-based population differentiation, GPA is a true generalist even at the level of a single clone, capable of shifting between highly divergent plant hosts (37–40). Previously, we reported that GPA Clone O, which descended from a single asexual female, successfully colonized nine hosts from five different plant families (41). Individuals that feed on one type of host can also survive and reproduce on distantly related host plants (41). These findings demonstrate that GPA can colonize diverse host plant species in the absence of genetic variation.

In this study, we investigated the RNA editing events in the transcriptome of GPA and found that GPA was capable of diverse editing events. Many RNA editing sites were responsive to host change. The GPA genome encoded two copies of *ADAR*s, which were responsible for A-to-I editing, while lacking AID/APOBEC deaminases. RNAi of each of the *ADAR*s resulted in a significant reduction in A-to-I editing levels and led to a reduced survival rate of GPA on host plants. These findings demonstrate that RNA editing fine-tunes transcriptomic plasticity and enables rapid, reversible adaptation to environmental change, even in the absence of genetic variation.

## Results

### RNA single-nucleotide variants in *M. persicae* vary according to host plants

Previously, we established stable colonies on nine plant species using colonies of *M. persicae* Clone O that derived from a single female. RNA-seq analysis of *M. persicae* on these hosts revealed host-specific patterns of coordinated gene expression (37). Remarkably, numerous single-nucleotide variants (SNVs) were detected in the RNA-seq reads aligned to the chromosome-level genome of Clone O (Fig. 1A).

**Fig. 1.**
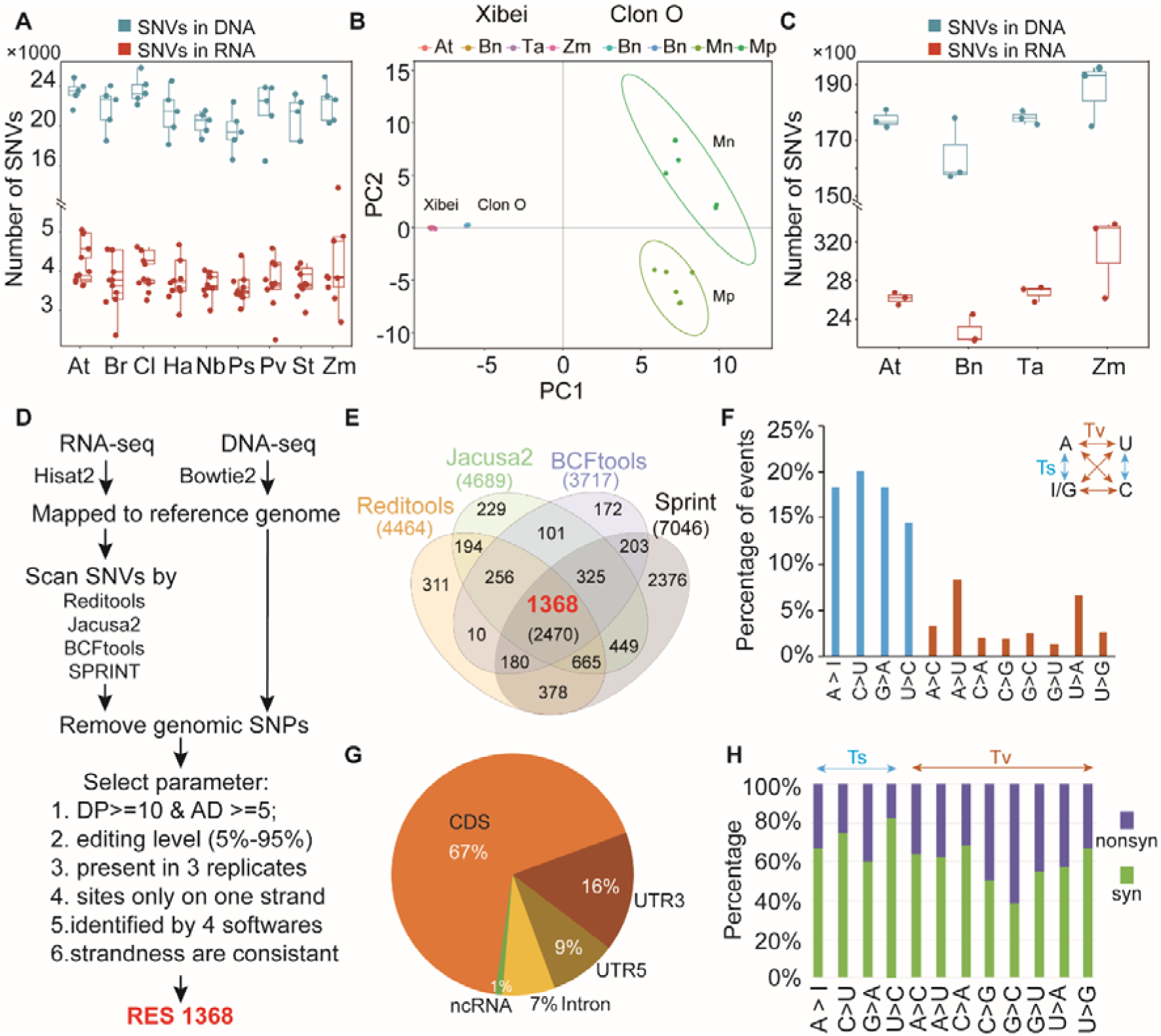
Mismatches between RNA and DNA are derived from RNA editing. (A) Number of SNVs detected in RNA and DNA of *M. persicae* Clone O across nine different host plants. RNA-seq data were collected from 2017 to 2018 and published in Chen *et al*. (2020) (41). DNA-seq data from 2016 and 2018 were published in Mathers *et al*. (2017, 2019) (37,42). Host plants include *Brassica rapa* (Br), *Arabidopsis thaliana* (At), *Nicotiana benthamiana* (Nb), *Solanum tuberosum* (St), *Chrysanthemum indicum* (Ci), *Helianthus annuus* (Ha), *Pisum sativum* (Ps), *Phaseolus vulgaris* (Pv), and *Zea mays* (Zm). (B) Single-female–derived GPA lines on different hosts (Clone O and Xibei) clustered tightly together, whereas the Mp and Mn geographic lineages were more divergent. Principal component analysis (PCA) of *M. persicae* colonies based on DNA-seq–derived SNVs. DNA-seq data for Clone O were obtained from Mathers *et al*. (2017, 2019), for the Xibei clone from Wu *et al*. (2024), and for Mp and Mn from Singh *et al*. (2021). (C) Number of SNVs detected in RNA and DNA of the Xibei clone across four different host plants: Bn, At, Ta, and Zm. (D) Workflow for identifying RNA editing sites. Strand-specific RNA-seq data were aligned to the *M. persicae* reference genome using HISAT2, and DNA-seq data were aligned using Bowtie2. SNVs in RNA were independently identified using Reditools, JACUSA2, BCFtools, and SPRINT. SNVs also present in the DNA were excluded. After applying six filtering criteria, 1,368 high-confidence RES were identified. DP (depth) indicates the total number of reads (edited and unedited), and AD (allele depth) refers to the number of edited reads. (E) Overlap of RES detected by the four tools: Reditools (46), JACUSA2 (47), BCFtools (48), and SPRINT (49). (F) Distribution of the 12 possible nucleotide substitution types among total RES. Transition events were more frequent than transversions. (G) Genomic features associated with RES. (H) Proportion of nonsynonymous (nonysn) and synonymous (syn) changes among RES located in coding sequences (CDS).

To investigate the origin of these SNVs, we compared the SNVs to those identified in DNA-seq data from aphids maintained on *Brassica rapa* (Br) before (2016) and after (2018) the nine-host experiment conducted in 2017 (37, 42). 78.7%–88.4% of SNVs in the RNA-seq corresponded to genomic mutations (Figure 1A). After subtracting the genomic SNVs, a significant number of SNVs remained in RNA-seq data from aphids on Br and on the other eight hosts (Figure 1A), indicating the presence of nucleotide mismatches in RNA relative to genomic DNA of *M. persicae*.

To elaborate on the occurrence of the RNA-specific SNVs in this species, we established stable colonies of *M. persicae* Clone Xibei (descended from a single female) on four different hosts, Bn (*B. napus*), At (*Arabidopsis thaliana*), Ta (*Triticum aestivum*), and Zm (*Zea mays*). Those aphid lines were maintained on corresponding plants for more than a year. Three aphid samples on each host were harvested for RNA-seq, and two additional samples for DNA-seq. Reads were aligned to the reference genome of Clone O. PCA of SNVs revealed that lines of Clone Xibei maintained on different host plants clustered closely (Fig. 1B), indicating that their genomes remained largely identical. Similarly, lines of Clone O, which were maintained on *B. napus* for over two years, exhibited few differences (Fig. 1B). After excluding DNA-based SNVs, thousands of RNA-specific SNVs were detected across the lines (Fig. 1C). This pattern was consistent across both Clone O and Clone Xibei (Fig. 1A and 1C), highlighting the presence of RNA-specific nucleotide variations in aphid lines of the same clone maintained on distinct host plants.

### RNA editing in *M. persicae* is diverse

SNVs in RNA that are independent of genomic variation may arise from RNA editing (43–45). To explore RNA editing in *M. persicae*, we employed four bioinformatic tools—REDItools2 (46), JACUSA2 (47), BCFtools (48), and SPRINT (49)—to identify RNA-editing sites (RES) by comparing RNA-seq data from Xibei clones with corresponding DNA-seq data. Strand-specific RNA-seq reads were aligned to the reference genome, and single-nucleotide mismatches between RNA and DNA were detected. SNVs present in the DNA samples were excluded to eliminate potential genomic mutations that could mimic RNA editing.

To ensure high-confidence identification of RNA editing events, we applied stringent filtering criteria (Fig. 1D). Candidate sites were required to (i) contain at least 10 uniquely mapped reads, including ≥5 edited reads, (ii) show an editing level between 5% and 95%, (iii) be consistently detected across all three biological replicates, and (iv) be called by all four prediction tools. Using these criteria, we identified 1,368 high-confidence RNA editing sites (Fig. 1E; Dataset S1). Of these, 951 sites (69.5%) were located in single-copy genes, 21.5% mapped to multi-copy gene regions, and the remaining 9.0% were found in intergenic regions (Fig. S1).

The identified RNA editing events were then categorized into transitions and transversions based on the type of nucleotide change (Fig. 1F). Transition events accounted for 71.6% of all editing sites and included A-to-I (18.4%), C-to-U (20.1%), G-to-A (18.4%), and U-to-C (14.6%) (Fig. 1F). Transversions comprised the remaining 28.4%, with A-to-U (8.6%) and U-to-A (6.7%) being the most frequent, while other transversion types occurred at lower frequencies, each representing less than 11% of the total (Fig. 1F). Independent analyses using the four software tools yielded highly consistent substitution patterns and relative frequencies (Fig. S2). While A-to-I editing is a common editing event in many organisms, the prevalence of other types of RNA editing in *M. persicae* is notable and suggests a distinct editing repertoire compared to other species (14, 17, 19, 24).

Analysis of the genomic distribution of RES revealed significant differences between editing types. Most RNA editing sites were located in coding sequences (CDS), and the impact of editing varied between transitions and transversions (Fig. 1G). Within the CDS, both transition and transversion edits generated a substantial number of nonsynonymous substitutions (Fig. 1H). The RNA editing profiles observed in *M. persicae* Clone O were further supported by high-depth RNA-seq data (37), confirming the robustness of the identified editing patterns (Fig. S3). Functional enrichment analysis revealed that CDS-resident RNA editing events in *M*. *persicae* were associated with cell adhesion and tissue morphogenesis, phosphorylation-dependent signal transduction, synaptic and neuromuscular structural organization, and extracellular matrix function (Fig. S4), suggesting that RNA editing may act as a key regulatory mechanism modulating developmental plasticity and host adaptation in *M. persicae*.

### RES are differentially edited in GPA on different host plants

To investigate the potential role of RNA editing in host adaptation by *M. persicae*, we identified RES that were differentially edited (DE) in response to different host plants. Editing levels for RES were obtained by Reditools and compared between lines of Clone Xibei maintained on four host plants, At, Zm, Ta, Bn. A total of 154 RESs were found to be DE among these hosts (Turkey test, log2FC ≥ 2, *P* < 0.05; Fig. S5A, Dataset S2). Of these, 53 RES were DE between aphids on At and Bn, 54 between those on Ta and Bn, and 99 between Zm and Bn (Fig. S5A, Dataset S2). A heatmap of editing levels indicated that RNA editing events were host-responsive, suggesting a potential role for RNA editing in enabling *M. persicae* to colonize different plant species (Fig. 2A).

**Fig. 2.**
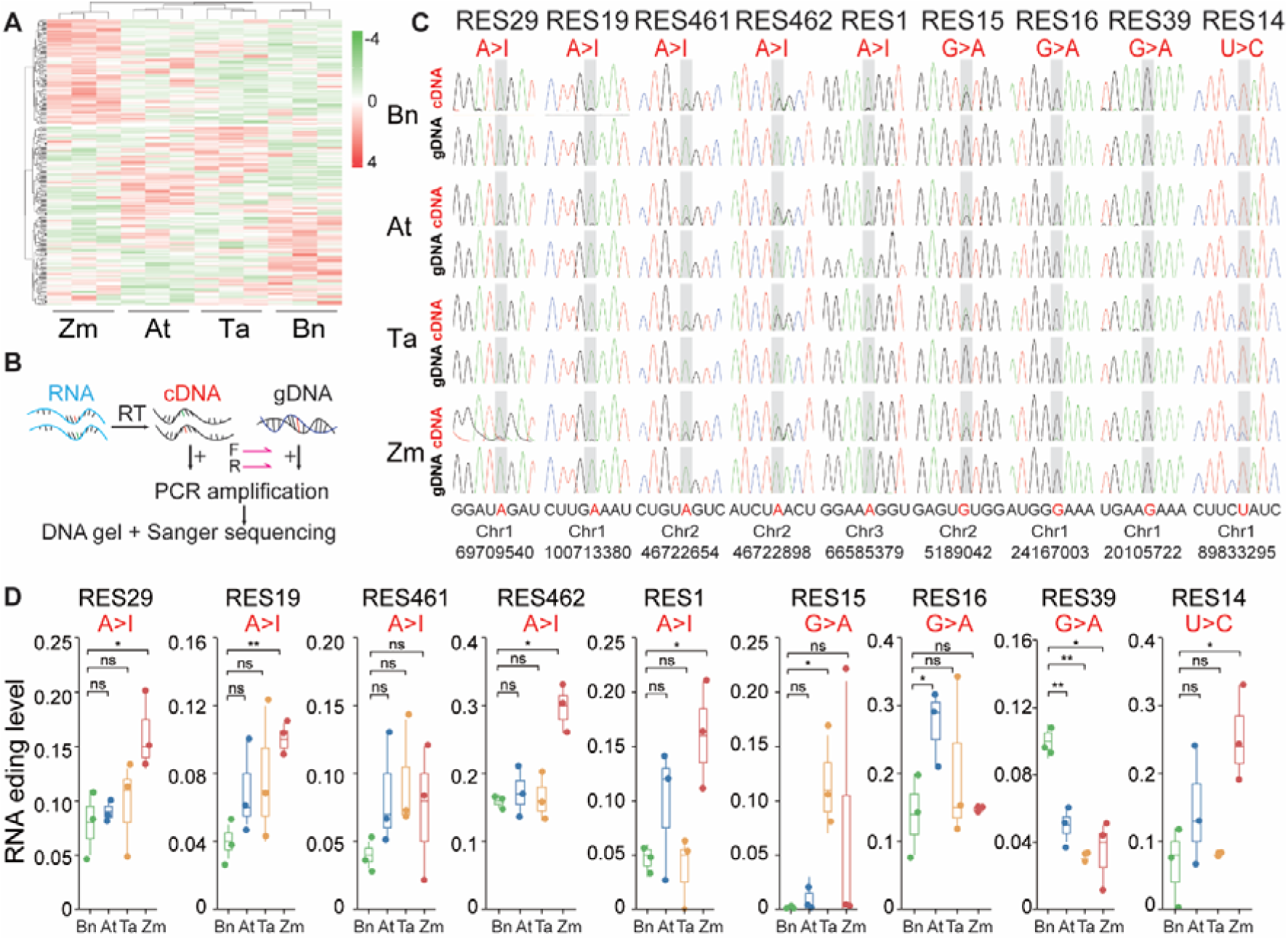
RES differentially edited in *M. persicae* on divergent plant hosts. (A) Heatmap of differentially edited RES across host plant treatments. (B) Workflow for RES validation. Forward and reverse primers were designed to amplify each target RES with 150–200 bp flanking sequences. PCR was performed using cDNA and gDNA as templates, followed by Sanger sequencing of the products. (C) Representative nucleotide sequences of RES in cDNA and gDNA from aphids reared on four host plants. Each panel is labeled with the corresponding RES identifier and editing type. Edited sites are highlighted in gray. In cDNA chromatograms, black peaks with gray highlights represent inosines (I) that were detected as guanosines (G) in Sanger sequencing. Genomic coordinates are indicated below the panels. (D) Editing levels of selected RES in aphids on different host plants. Colored dots represent RNA editing levels of individual samples for each host, and boxplots show the data distribution within each host. Statistical significance is indicated as \**P* < 0.05, \*\**P* < 0.01, \*\*\**P* < 0.001, and n.s. denotes not significant (*P* > 0.05), based on Wilcoxon rank-sum test.

To validate these RES, PCR amplification and sequencing were performed using specific primers targeting the identified RES (Fig. 2B). Among the 154 RESs (Dataset S2), 71 sites were amenable for primer design (Table S1). Primers covered RES sites were consistently and specifically amplified from cDNA and gDNA samples of aphids on the four host plants (Fig. S5B). For verification, we used biological samples different from those employed in the RNA-seq analysis to ensure the reliability of RES across lines. Sequence analysis revealed that nine RES were present in PCR products derived from cDNA but absent from gDNA (Fig. 2C). The nine RES included five A-to-I sites, three G-to-A sites, and one U-to-C site (Fig. 2C), which were repeatedly validated across multiple independent samples. TPM of editing levels at these sites varied significantly across lines reared on different host plants (Fig. 2D), further supporting the hypothesis that RNA editing contributes to host adaptation in GPA.

### Rapid response of RES editing levels correlates with gene expression to host change

To determine whether RNA editing levels in GPA respond to different host plants, we developed a RES-specific qPCR approach to quantify the editing levels of specific RNA editing sites modified from allelic-specific PCR (50) and mismatch primer design (51–53) (Fig. 3A). This method was designed to distinguish between edited and unedited transcripts by placing the edited or unedited nucleotide at the 3′ end of the forward primer and introducing an additional mismatch 1–2 bases upstream. These primers, combined with a common reverse primer, allowed for specific amplification of edited versus unedited cDNA templates (Fig. S6A).

**Fig. 3.**
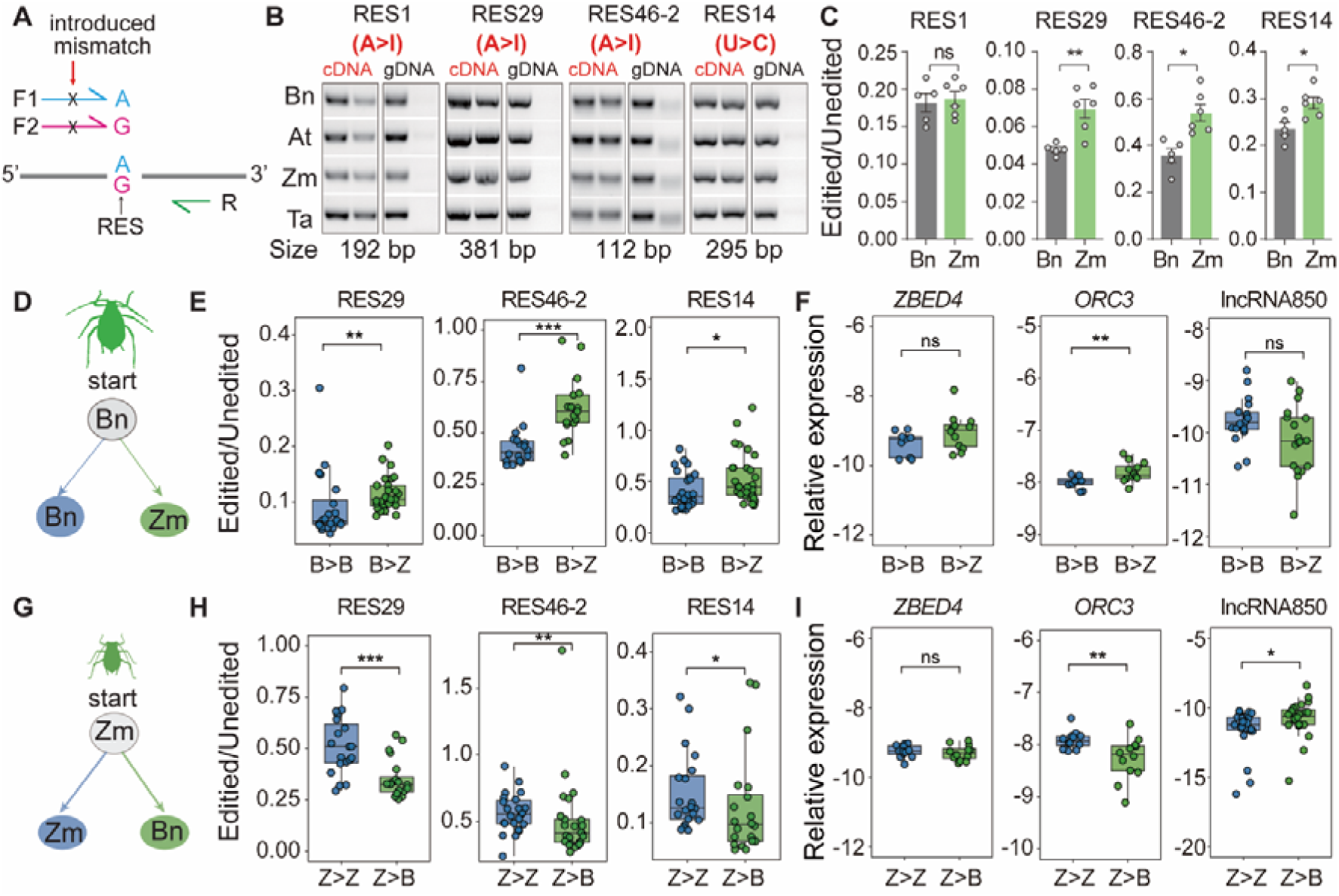
Rapid response of RES editing levels and gene expression to host change. (A) Design of RES-qPCR. The 3′ end of each forward primer targets the edited or unedited nucleotide, with an additional mismatch introduced 1–2 nucleotides upstream. Each forward primer is paired with a common reverse primer to selectively amplify edited or unedited transcripts. (B) RES-qPCR selectively amplifies both edited and unedited transcripts from cDNA, but only unedited transcripts from gDNA. (C) Ratio of edited to unedited transcripts in *M. persicae* reared on Bn and Zm. (D, G) Schematics of host-switch experiments. Aphids reared on Bn were transferred either to Zm (B>Z) or to Bn (B>B), and aphids reared on Zm were transferred either to Bn (Z>B) or to Zm (Z>Z). (E, H) Ratio of edited to unedited transcripts following host transfer from Bn to Zm and from Zm to Bn, respectively. (F, I) Expression levels of genes containing RES after host transfer from Bn to Zm and from Zm to Bn, respectively. In Figures C, E, and H, RNA editing levels were quantified by qPCR. For each gene, the total transcript abundance was used as an internal reference. ΔCt values for the edited and unedited transcripts were calculated relative to this reference and subsequently converted to relative expression levels (2^^–ΔCt^). The RNA editing level was then expressed as the ratio of 2^^–ΔCt(edited)^ to 2^^–ΔCt(unedited)^, yielding results consistent with those normalized using conventional housekeeping genes. In Figure F and I, relative expression of genes containing candidate editing sites was calculated using the 2^^–ΔCt^ method and normalized to the *EF1*α gene. RES29 corresponds to *ZBED4*, RES46-2 to *ORC3*, and RES14 to long non-coding RNA 850 (lncRNA850). In Figure C, E, F, I, and J, each data point (white dot with a black outline) represents a ratio or expression level, and bar graphs show the distribution across treatments. Each treatment includes n = 15 aphids. Statistical significance is indicated as \**P* < 0.05, ** *P* < 0.01, *** *P* < 0.001, and n.s. denotes not significant (*P* > 0.05), determined by Wilcoxon rank-sum test.

We first validated primer specificity by showing that they amplified edited and unedited cDNA, but not gDNA (Fig. 3B). Sanger sequencing confirmed that the PCR products contained the expected edits and primer-introduced mismatches (Fig. S6B). Edited and unedited primers showed similar amplification efficiencies on cDNA, but amplification of gDNA with edited primers was strongly suppressed due to the 3′-end mismatch (ΔCt = 5.36–9.05) (Fig. S6C), which is consistent with previous reports (53). Using this RES-qPCR method, we quantified three A-to-I sites (RES1, RES29, RES46-2) and one U-to-C site (RES14) (Fig. 3B). Editing at three of these sites—RES29, RES46-2, and RES14—varied with host plant, showing higher levels in GPA feeding on Zm than on Bn (Fig. 3C).

To further test whether RNA editing responds dynamically to different host plants, we performed reciprocal host-switch experiments (Fig. 3D, 3G). Third-instar GPAs reared on Bn were transferred to either Bn (control) or Zm. After 48 hours, editing at RES29, RES46-2, and RES14 was significantly increased in aphids switched from Bn to Zm (Fig. 3E). RES14 (a U-to-C event) is located in the 5′ UTR of a lncRNA (MYZPE13164_O_EIv2.1_0099850, hereafter lncRNA850), whereas RES46-2 (an A-to-I event) resides in the 3′ UTR of the *ORC3* gene. Consistent with the induction of editing, lncRNA850 expression decreased, and expressions of *ZBED4* and *ORC3* were increased in Bn-to-Zm aphids (Fig. 3F). The opposite trend was observed in the reciprocal switch. When GPAs were transferred from Zm to Bn (Fig. 3G), editing levels at RES29, RES46-2, and RES14 significantly decreased (Fig. 3H). Correspondingly, expressions of *ZBED4* and *ORC3* were downregulated, and lncRNA850 expression was upregulated in Zm-to-Bn aphids (Fig. 3I).

Together, these results suggest that RNA editing in GPA is responsive to host plants and may play a role in host adaptation, likely through site-specific RNA editing events affecting both coding and non-coding transcript features.

### Identification and domain architecture of RNA-editing deaminases in GPA

In eukaryotes, the RNA editing mechanisms remain largely unknown. Among the best-characterized events are transition edits such as A-to-I and C-to-U, mediated by deamination reactions. Few U-to-C and G-to-A events resulting from transamination reactions (33, 48) have been reported, but the enzymes responsible for these are still unknown. Therefore, we focused on identifying deaminases in *M. persicae*.

Using the Pfam database, we surveyed the deaminase superfamily (Pfam clan CL0109), which includes 33 domain families (Dataset S3), along with two additional domains—dsRNA-binding (dsRB) and Z-DNA/RNA-binding—known from mammalian ADARs. We searched for all 35 domains in the GPA Pfam annotations (Dataset S4) and identified the presence of A_deamin and dsRB domains, but not the Z-DNA/RNA-binding domain (Dataset S3), suggesting that GPA ADARs contain A_deamin and dsRB domains. Three proteins with A_deamin domains were found: two also contained dsRB domains and were annotated as ADAR-like based on NCBI homology, while the third was annotated as an ADAT (adenosine deaminase acting on tRNA) (Dataset S4). No domains related to AID/APOBEC deaminases were detected in GPA or other arthropod genomes examined.

ADARs and ADATs both catalyze A-to-I RNA editing, but act on distinct substrates: ADARs on double-stranded RNAs and ADATs on tRNAs. Comparative analysis of 22 arthropod genomes revealed that the ADAT gene is present as a single copy in most species (Dataset S3), with multiple copies detected only in a few cases. In contrast, ADAR genes occur as multiple copies in several species, and some copies lack canonical dsRB domains (Fig. 4A). These findings indicate that ADAR genes have undergone lineage-specific diversification in arthropods, which may lead to differences among species in RNA substrate recognition or post-transcriptional regulatory functions.

**Fig. 4.**
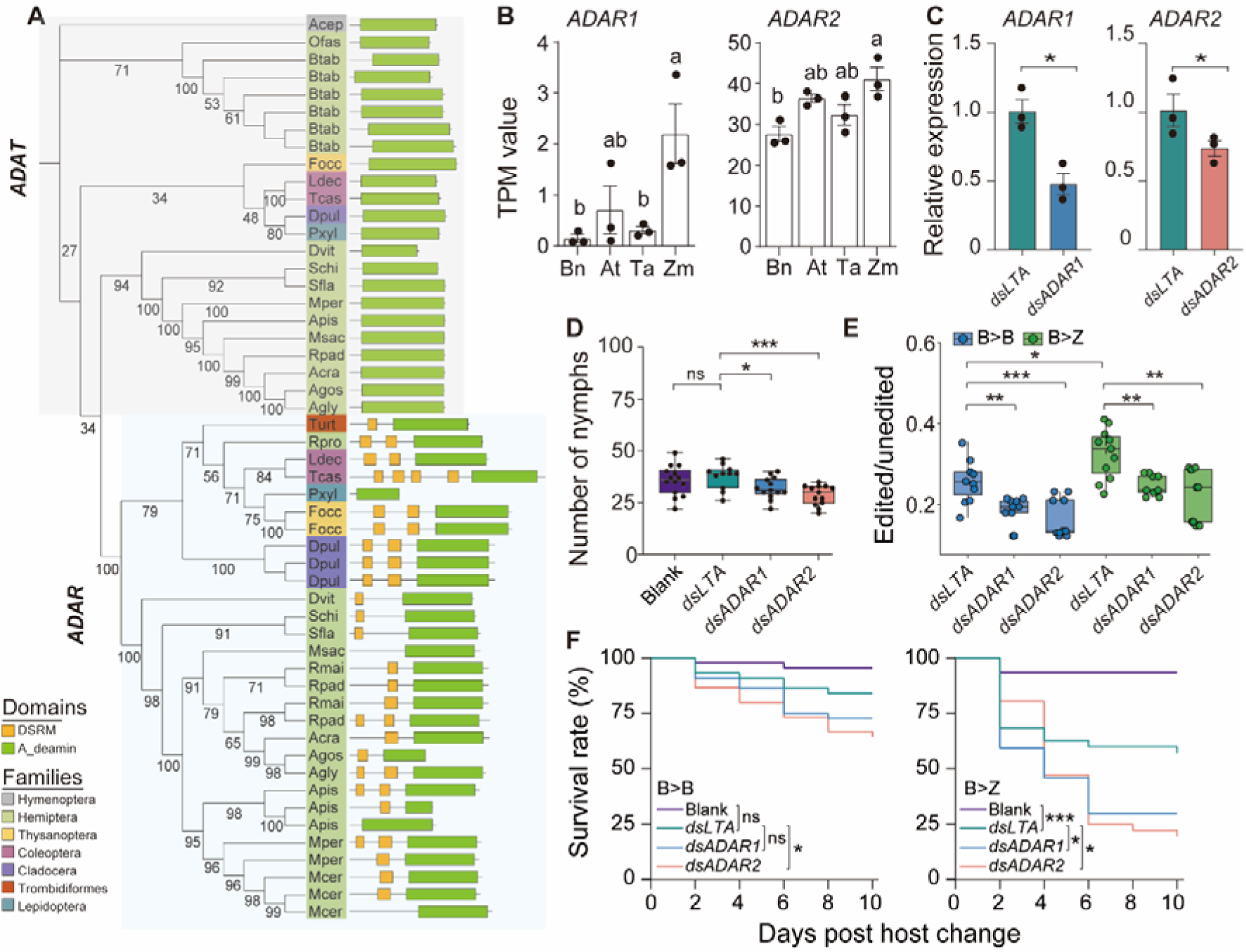
ADARs regulate RNA editing and contribute to *M*. *persicae* host adaptation. (A) Phylogenetic tree of ADARs and ADAT. *Tetranychus urticae* was used as the outgroup. Bootstrap values (%) are indicated at the nodes. Shaded colors indicate insect orders. Branch labels show species abbreviations (e.g., Mper = *M. persicae*; full list in Table S3). Functional domains are shown to the right: yellow boxes = dsrm (double-stranded RNA-binding) domain, green bars = adenosine deaminase domain. The large gray and blue shaded regions highlight the ADAT and ADAR clades, respectively. (B) TPM of *ADAR1*, *ADAR2* in *M. persicae* reared on different host plants. Different lowercase letters indicate significant differences (one-way ANOVA with Tukey’s test). (C–D) RNAi knockdown of *ADAR1* and *ADAR2* by dsRNA feeding reduced transcript levels (normalized to *MpEF1α*, mean ± SEM, n = 3), decreased fecundity (each dot = 1 aphid, n = 12–15; blank: 30% sucrose, *dsLTA*: unrelated dsRNA). (E) Silencing *ADAR1* or *ADAR2* reduced RES46-2 editing levels on Bn and blocked the rapid upregulation observed after transfer to Zm (mean ± SEM, n = 10). (F) Silencing *ADAR1* or *ADAR2* reduced survival on the new host (surviving aphids counted every two days, n = 10–12). Statistical significance: Wilcoxon rank-sum test for transcript levels, fecundity and editing levels, log-rank test for survival; \*\*\**P* < 0.001, \*\**P* < 0.01, \**P* < 0.05, n.s = not significant.

### *ADARs* are essential for GPA host adaptation

In GPA, expression of *ADAR* genes varied significantly depending on the host plant (Figure 4B), and their expression was increased in response to changed host plants (SI Figure 7), implicating their potential roles in RNA editing-mediated adaptation to different hosts. To directly assess the functional contribution of ADAR genes to host adaptation, we silenced *ADAR1* and *ADAR2* in aphids originally maintained on Bn via dsRNA feeding (Fig. 4C). Knockdown of either gene significantly reduced aphid fecundity compared with aphids fed a blank treatment or *dsLTA* control (Fig. 4D). These findings, together with the host-dependent expression patterns of *ADAR*s, indicate that *ADAR*s are required for GPA fitness and likely play an essential role in host plant adaptation.

To test the impact of *ADAR*s in host adaptation, we performed host-change experiments by transferring RNAi-treated and control aphids from Bn to Zm. In *ADAR1*- and *ADAR2*-silenced aphids on Bn, the editing level at RES46-2—a site of A-to-I editing—was decreased relative to control. After 24 hours on Zm, control aphids exhibited upregulation of both RES46-2 editing (Figure 4E), which was abolished in RNAi aphids (Figure 4E). As a result of this impaired response, *ADAR1*-and *ADAR2*-RNAi aphids experienced significantly higher mortality following transfer to Zm compared to non-silenced aphids (Figure 4F).

These results indicate that RES46-2 editing is regulated by ADARs in response to host plant changes. Disruption of the *ADAR*-mediated pathway is associated with lower GPA survival on a novel host, underscoring the role of RNA editing in host adaptation.

### Conserved A-to-I RNA editing with species-specific functional divergence across aphid lineages

To investigate RNA editing in aphids, we analyzed editing events in two specialist species, *Rhopalosiphum padi* (BOA) and *Brevicoryne brassicae* (CA), together with the generalist GPA. Previously, we generated RNA-seq datasets for GPA, CA, and BOA subjected to host plant shifts (38) and corresponding genomic DNA sequences (PRJNA1344291). After filtering out genomic SNPs using Reditools, we identified 1270 RNA editing sites in GPA, 368 in CA, and 500 in BOA (Fig. 5A, Dataset S5).

**Fig. 5.**
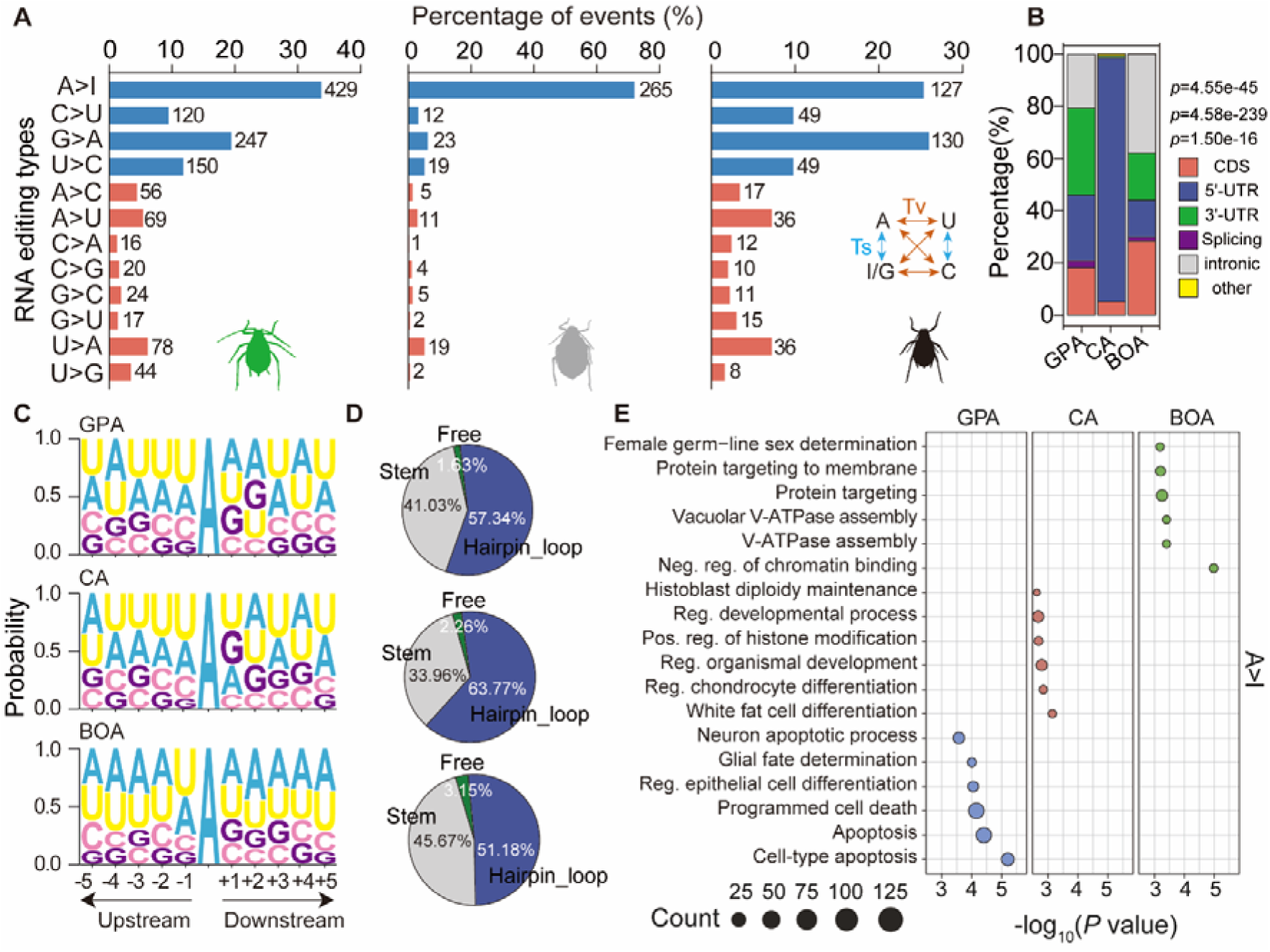
Comparative analysis reveals conserved and species-specific RNA editing patterns in aphids. (A) Distribution of the 12 types of RES in *M. persicae* (GPA, green), *B. brassicae* (CA, gray), and *R. padi* (BOA, black). The number of RNA variants for each nucleotide substitution type is shown to the right of each bar. A-to-I is the most prevalent type across all species. (B) Genomic distribution of A-to-I RES in the three aphid species. The stacked bar chart shows the percentage of editing events located in coding sequences (CDS), 5′-UTRs, 3′-UTRs, splice sites, intronic regions, and other regions. Significant differences in regional preference among species are indicated (*P* values from χ² tests). (C) Sequence conservation of A-to-I editing sites and nucleotide preferences in the flanking sequences (5 bp upstream and downstream). (D) Percentage of A-to-I RES located in different types of RNA secondary structure elements. Prediction of RNA secondary structures is based on 30-nt upstream and 30-nt downstream sequences surrounding the edited sites. (E) Gene Ontology (GO) enrichment analysis of genes harboring A-to-I RES. GO terms were detailed in Table S4. Enriched biological processes are plotted based on gene count (dot size) and significance (−log_10_ (*P value*)).

The genomic distribution of all identified editing sites also showed clear lineage specificity. The three species differed in editing preferences between untranslated regions (UTRs) and coding sequences (CDSs), and CDS edits contributed differently to synonymous and nonsynonymous substitutions (Fig. 5B). Across all three species, A-to-I was the predominant editing type, accounting for 33.8% in GPA, 72.0% in CA, and 25.4% in BOA. Notably, GPA and BOA exhibited a broader diversity of non–A-to-I editing events than CA, a pattern consistent with their wider host ranges (Fig. 5A).

Sequence context analysis revealed conserved motifs and characteristic RNA secondary structures surrounding A-to-I sites, which were preferentially located within stem–loop regions (Fig. 5C–D), supporting that A-to-I editing is a conserved and structurally guided feature of aphid transcriptomes. Despite this conservation, the functional targets of RNA editing differed among species (Fig. 5D, Fig. S8). In GPA, edited genes were enriched in neural and apoptotic pathways, including cell fate determination, differentiation, and programmed cell death. In BOA, editing events were concentrated in genes related to V-type ATPase assembly and chromatin regulation. In CA, edits were enriched in developmental and histone modification pathways (Fig. 5E, Table S4).

Overall, A-to-I RNA editing represents a conserved molecular mechanism in aphids, but its genomic distribution and functional targets are species-specific, reflecting potential differences in ecological adaptation and host utilization strategies.

## Discussion

In this study, we investigated parthenogenetic lines of GPA, a highly generalist aphid capable of colonizing over 1,000 plant species across more than 50 plant families (34). We show that extensive RNA editing in GPA generates substantial transcriptomic variation, potentially compensating for its limited genetic diversity. These editing events are diverse and, in many cases, strongly influenced by host plants, with editing levels shifting in response to different hosts. Notably, several RNA editing sites respond rapidly to host changes, correlating with altered expression of the corresponding transcripts.

RNA editing in GPA is highly dynamic. More than 150 editing sites differed among aphids feeding on four host plants, and several showed reversible shifts within 48 hours of host transfer. Editing occurred in both protein-coding and non-coding regions and was associated with host-responsive changes in transcript abundance. Of particular interest, we identified an A-to-I site (RES46-2) in the 3′UTR of *ORC3* and a U-to-C site (RES14) in the 5′UTR of a long noncoding RNA (lncRNA850). *ORC3* is a subunit of the origin recognition complex, a conserved multi-protein assembly that initiates DNA replication, and in insects it is implicated in replication licensing, cell cycle regulation, and developmental plasticity (54–58). These observations suggest that RNA editing may modulate transcript stability or translation in a host-specific manner, providing a mechanism for fine-tuning fundamental cellular processes in response to environmental change.

RNA editing likely underlies the transcriptional plasticity that allows GPA to thrive on diverse hosts. It confers several adaptive advantages: it enables rapid phenotypic plasticity without heritable genetic changes; allows environment-specific regulation of gene function through post-transcriptional modifications; and can modulate key physiological activities via nonsynonymous edits that alter protein function. For example, RES1 represents an A-to-I edit in the ion channel transcript *KCNB1* (Table S5), which encodes a voltage-gated potassium channel, suggesting that RNA editing may influence neural or physiological processes critical for host recognition, feeding, or detoxification.

RNA editing events in GPA can be classified into transitions and transversions, with transitions being the most common. GPA possesses two *ADAR* genes, each containing 1-2 double-stranded RNA-binding motifs, but lacks genes encoded for AID/APOBEC deaminases, which mediate other RNA editing types in many organisms (31). Knockdown of *ADAR* genes significantly reduced editing, decreased aphid fecundity, and impaired adaptation to novel hosts, demonstrating that RNA editing is essential for environmental acclimation in this parthenogenetic species. The enzymes responsible for less common editing types in GPA remain unknown, and understanding their origins may provide insights into the molecular basis of ecological generalism and host adaptation in insects.

A-to-I editing is the most common RNA editing type across metazoans, with other types observed less frequently. In GPA, however, RNA editing is exceptionally diverse, encompassing 12 types, many of which are rarely reported in other organisms (24). Among 22 metazoan species studied, A-to-I dominates, with other types largely absent except in the marine invertebrate *Trichoplax adhaerens*, which exhibits a broader editing repertoire (17, 24). Even in *Metopolophium dirhodum*, a close relative of GPA and a specialist on cereal plants, A-to-I remains predominant (17). In GPA, A-to-I edits account for only a fraction of the total, raising the intriguing question of whether the broader diversity of RNA editing types contributes to its polyphagy. Indeed, we observed a correlation between RNA-editing diversity and host range among aphids: GPA and BOA, which colonize multiple plant families, exhibit more non-A-to-I editing events than CA, a Brassicaceae specialist (59). Although A-to-I editing retains conserved biochemical features across aphids, its genomic distribution and functional consequences are species-specific, reflecting lineage-specific adaptation strategies.

In summary, GPA employs a diverse, dynamic, and host-responsive RNA editing system that overcomes the constraints of clonal reproduction and facilitates adaptation to a wide range of host plants. More broadly, these findings highlight RNA editing as a flexible and underappreciated driver of environmental adaptation in metazoans, particularly in species with limited genetic variation.

## Acknowledgments

This project is funded by the National Natural Science Foundation of China (project No. 32172392 to Y.C.), the National Key Research and Development Program of China (project No. 2023YFF1000703 to Y.C.), and supported by Hubei Hongshan Laboratory (project No. 2022hszd026 to Y.C.), the Startup Foundation for Advanced Talents at HZAU to Y.C., the Fundamental Research Funds for the Central Universities (Program No. 2022ZKPY003 to Y.C.), the Wuhan Yingcai Talent Program to Y.C., and grants from the National Science Fund for Distinguished Young Scholars (32325039 to S.J.).

## Author Contributions

Y.C., S.J., and G.W. conceived and designed the study, supervised the project, and finalized the manuscript. W.Y. performed the majority of the experimental work, and Y.C. and W.Y. jointly conducted the primary data analyses. J.W., Z.L., and P.P. contributed to the experimental setup and bioassays. Bioinformatic analyses were collaboratively carried out by Y.C., S.Z., W.Y., and R.H. P.Y. and Z.H. contributed to the execution of the research. The manuscript was drafted by Y.C. and W.Y., with W.Y. also managing the next-generation sequencing data. S.H., J.-C.S., G.W., S.J., and S.Z. provided academic consultation and contributed to manuscript revision. All authors reviewed the manuscript, provided feedback, and approved the final version.

## Data deposition

All data reported in this paper were deposited in the NCBI BioProject under project numbers PRJNA1211840 and PRJNA1344291. All data are available in the main text or the supplementary materials.

## Legends for Datasets S1 to S6

**Dataset S1: List of high-confidence RNA editing sites (RES) identified in *Myzus persicae***

Table 1: RES identified by REDItools2

Table 2: RES identified by JACUSA2

Table 3: RES identified by BCFtools

Table 4: RES identified by SPRINT

Table 5: RES identified by all four tools

### Dataset S2. Differentially edited RNA editing sites (DE-RES) in *Myzus persicae* in response to host plants

Table 1: DE-RES between At and Bn

Table 2: DE-RES between Ta and Bn

Table 3: DE-RES between Zm and Bn

**Dataset S3: Distribution of deaminase-related Pfam domains across 21 arthropod species**

**Dataset S4: Pfam domain annotations of ADAR-related proteins in *Myzus persicae***

**Dataset S5: RES identified in GPA, CA, and BOA under host-shift treatments.**

Table 1: RES identified in *M. persicae* (GPA) across four host-shift conditions (B > B, B > Z, Z > Z, Z > B)

Table 2: RES identified in *B. brassicae* (CA) under two host-shift conditions (B > B, B > Z)

Table 3: RES identified in *R. padi* (BOA) under two host-shift conditions (Z > Z, Z > B)

## Materials and methods

### Aphid rearing

*M. persicae* (GPA) Clone Xibei was obtained from a stock culture maintained on cabbage (*B. oleracea*) in Prof. Tongxian Liu’s laboratory at Northwest A&F University (Yangling, Shaanxi). This clone was subsequently propagated into lines maintained on four host species: *Z. mays* (Zm) B73, *B. napus* (Bn) Zhongshuang 11, *A. thaliana* (At), and *T. aestivum* (Ta). The GPA was initially reared on Bn for 2–3 weeks, after which a single parthenogenesis female was transferred to Zm, At, and Ta to establish stable parthenogenetic lines. The four aphid lines were maintained at 24 ± 1°C, 60 ± 10% relative humidity, and a 16 h light/8 h dark photoperiod for over 2 years.

For each GPA line, five samples were collected: three for RNA sequencing and the remaining two for DNA sequencing. Each sample comprised 30 age-synchronized adults.

### RNA extraction and RNA sequencing

Aphid samples for RNA-seq were ground to a fine powder with sterilized stainless-steel beads in a TissueLyser II (Jingxin, Shanghai, China). Total RNA was isolated and purified from the samples using TRIzol® Reagent (Invitrogen,15596026CN, USA) in conjunction with VAHTS® mRNA Capture Beads (Vazyme, N403, Nanjing, China). RNA concentration and purity were quantified using a NanoDrop 2000 spectrophotometer (Thermo Fisher Scientific, USA) and Qubit 2.0 Fluorometer (Thermo Scientific, USA). RNA integrity was subsequently evaluated using the Agilent 2100 Bioanalyzer (Agilent Technologies, USA).

Strand-specific libraries were constructed following the protocol provided with the VAHTS® Universal V6 RNA-seq Library Prep Kit (Vazyme, NR604, Nanjing, China). Sequencing was then performed on the DNBSEQ-T7 platform (OEbiotech, Shanghai, China), generating paired-end reads of 150 bp.

### DNA extraction and DNA sequencing

Genomic DNA of each GPA line, was extracted using the cetyltrimethylammonium bromide (CTAB) method and further purified with the Blood and Cell Culture DNA Midi Kit (QIAGEN, 13343, Germany).

Library construction and genome sequencing were performed by Novogene (Beijing, China) according to the manufacturer’s protocol. Sequencing libraries were generated using the NEBNext Ultra DNA Library Prep Kit (Illumina, E7645S, USA). Sequencing was then carried out on the Illumina NovaSeq 6000 platform (Illumina, USA), with paired-end reads of 2 × 150 bp for each sample.

### SNP Calling by GATK

To minimize false-positive SNVs introduced by sequencing errors or mapping bias, we implemented stringent multi-step filtering and cross-validation. In addition to standard GATK filtering (QUAL, QD, FS, SOR, and cluster thresholds), we verified variant calling accuracy by benchmarking with simulated datasets of known variants, ensuring tool performance under our sequencing depth and read length conditions. Variants supported only by a single aligner or by strand-biased reads were discarded. This conservative pipeline reduces spurious SNVs and increases confidence in downstream RNA editing detection.

We used the published genome sequencing data of the *M. persicae* Clone O strain (PRJEB11304, PRJNA613055) and previous transcriptome sequencing data of GPA colonized on nine host plants (PRJEB24317, PRJEB11304, PRJNA532419) (1–3), for subsequent analysis. DNA-seq and RNA-seq data were aligned to the *M. persicae* Clone O genome (4) using BWA (version 0.7.17-r1188) (5) and HISATS2 (6) (version 2.2.1), respectively. The alignment results were converted to BAM format using SAMtools (version 1.18) (7). The BAM files were then sorted and indexed using SortSam, and potential PCR duplicates were marked using MarkDuplicates both from the Picard Toolkit (version 3.0) (broadinstitute.github.io/picard/). Additional indexing was performed with SAMtools to facilitate downstream analyses. Single nucleotide polymorphisms (SNPs) were identified using HaplotypeCaller from the GATK (version 4.1.8.1) tool (8), and multi-dimensional filtering was applied to retain high-quality variants: (a) quality score (QUAL) below 30 were discarded; (b) quality-by-depth ratio (QD) below 2 were excluded to avoid variants influenced by abnormal sequencing depth; (c) significant strand bias, as indicated by Fisher Strand Bias (FS) and Symmetric Odds Ratio (SOR) values exceeding 60 and 4, respectively, were removed; (d) clusters of more than three variants within a 10-base window were identified and excluded as potential false positives; and (e) missing values were considered non-passing variants. Then, VCF files from all samples were merged using BCFtools (version 1.8) (9). To eliminate genomic SNPs from the transcriptomic data, DNA variant sites were excluded using VCFtools (version 0.1.16) (10), resulting in a final set of transcriptome-specific SNPs.

### Ortholog analysis

Protein sequences of GPA were analyzed using OrthoFinder (v2.5.4) (11) with the parameters “-M msa -t 30 -a 30”, which identified 23,912 genes in *M. persicae*, including 8,109 single-copy genes (33.91%) and 15,803 multi-copy genes (66.09%).

### GO term enrichment analysis

GO term enrichment analyses using the clusterProfiler package (version 3.18.1) (12). The input for these analyses was a series of defined DEG IDs.

### Identification of RNA-editing sites

To enhance the robustness of RNA editing site (RES) detection, we employed four independent tools (REDItools2, JACUSA2, BCFtools, and SPRINT) and retained only consensus sites consistently identified by all tools. To further minimize potential mapping artifacts, candidates located in paralogous or repetitive genomic regions annotated by RepeatMasker were excluded. To ensure reproducibility, only sites present in all three biological replicates were considered. RES identification was based on paired transcriptome and genome resequencing data generated in this study, following the principle that target sites must be homozygous in genomic DNA (gDNA) and exhibit consistent nucleotide mismatches between RNA and DNA alignments (13). Based on this principle, two sequential rounds of RES identification and filtering were performed. In the first round, SNP detection was conducted on both RNA-seq and DNA-seq data to identify transcription-derived variants. Raw RNA-seq and DNA-seq reads were quality-controlled using fastp (v0.23.4) (14) and aligned to the reference genome as previously described. RNA-seq editing sites were independently identified using four bioinformatics tools: REDItools2 (v2.0) (15) with the REDItoolDnaRna.py script (-S -mrl 30 -q 20 -bq 30 -men 1 -T 1 -Tv 0.7 -N - B); JACUSA2 (v2.04) (16) in call-1 -a D -p 20 -f V mode; BCFtools (v1.10) (17) using mpileup and call functions (--max-depth 10000 -q 20 -Q 20); and SPRINT (v0.1.8) (18) (-cd 200 -csrg 5). SNP detection for DNA-seq data was performed as described above. RES identified from RNA-seq data were then filtered against gDNA SNPs to exclude sites originating from genomic variation. In the second round of filtering, high-confidence RES were defined using the following criteria: (a) Coverage depth (DP) ≥ 10 and supporting reads for the edited base (AD) ≥ 5; (b) Editing level between 5% and 95%; (c) Presence in all three biological replicates; (d) Occurrence on a single strand only; (e) Identification by all four bioinformatics tools; (f) Consistency of editing results between forward and reverse strands.

### Genomic feature of RNA editing sites

RNA editing sites (RES) were annotated using SnpEff (v5.2) with the parameters *-canon -no-upstream -no-downstream -hgvs* (19), restricting annotation to canonical transcripts and avoiding ambiguous intergenic assignments. Annotation incorporated aphid-specific gene models from Liu *et al*. (4). Because SnpEff assumes eukaryotic transcript structures, putative prokaryotic-like RNA modification features were excluded. For each RES, genomic context—including UTRs, introns, exons, and lncRNAs—was determined based on the *M. persicae* genome annotation (4). Additionally, the numbers of nonsynonymous and synonymous mutations were quantified to assess their potential impact on protein-coding sequences. To further explore sequence features and RNA structures, 30 nucleotides upstream and downstream of each A-to-I RNA editing site were extracted. WebLogo 3 (20) was used to evaluate and visualize sequence preferences surrounding editing sites, while RNAFold (21) was employed to predict the secondary structures of RNA sequences containing edited A sites.

### Differential editing (DE) RES analysis

To assess whether RNA editing levels were significantly influenced by host plant species, we performed linear regression analyses using R. Editing levels (dependent variable: value) were modeled as a function of host identity (independent variable: hosts) for each editing site individually.

For each RNA editing site, we applied a linear model: *lm* (*value* ∼ *hosts*). Here, *value* represents the editing level, and hosts is a categorical variable representing the host plant. The regression was conducted using the dplyr package in R by grouping the Dataset by editing site and applying the linear model across each group.

To evaluate the effect of the host on editing level, we extracted the *p*-value associated with the *hosts* coefficient from the model summary. This corresponds to an ANOVA F-test assessing whether host species explains a significant proportion of the variance in editing level. The analysis was performed using basic R functions, and the resulting *p*-values were used to determine host-dependent RNA editing events.

To define DE RES, we applied two criteria: a statistically significant difference in editing levels across hosts (*P* < 0.05), and a fold-change in editing level of ≥2 between at least two host conditions.

### Verification of the candidate RES by Sanger sequencing

To validate whether the candidate sites are edited in *M. persicae*, primers were designed with 150-200 bp flanking sequences upstream and downstream of the candidate sites (Figure 2B). PCR amplification was performed using cDNA and genomic DNA (gDNA) as templates, followed by Sanger sequencing of the amplified products. For cDNA synthesis, 1 µg of total RNA was reverse transcribed using the PrimeScript™ RT Reagent Kit with gDNA Eraser Kit (TaKaRa, RR047A, Japan) according to the manufacturer’s instructions. The PCR reaction was typically conducted in a 25 µL volume, containing Phanta® Max Super-Fidelity DNA Polymerase (Vazyme, P505, Nanjing, China), 100 ng of cDNA (or 10 ng of gDNA) template, and 10 µM each of forward and reverse primers. The PCR program was set as follows: 95°C for 3 minutes for initial denaturation, followed by 38 cycles of 95°C for 15 seconds, 57°C for 15 seconds, and 72°C for 30 seconds, with a final extension at 72°C for 5 minutes. Primers were synthesized by Shanghai Sangon Biotech Co., Ltd., and Sanger sequencing was performed by Wuhan Quintarabio Biotech Co., Ltd. The presence of RNA editing was visualized using SnapGene.

### RES-specific qPCR

To assess the variation in editing levels of candidate sites across different hosts, we propose an efficient and convenient quantification method—RES-specific qPCR (RES-qPCR). This method exploits the mismatch effect between the template and primers to influence the DNA polymerase amplification efficiency, leading to changes in fluorescence signals, thereby enabling the quantification of different transcript variants. The primer design strategy is adapted from genome-wide allele-specific SNP detection methods (22, 23). For each candidate editing site, a set of primers is designed, including two specific editing primers and one universal primer (Figure 3A). When the specific editing primer serves as the forward primer, the universal primer is designed as the reverse primer, and vice versa. The two specific editing primers differ only at the first base of the 3’ end, allowing effective discrimination between edited and unedited sites. To further enhance primer specificity, a second mismatch is introduced at the 3’ end of the specific editing primers, either at the first or second position upstream. The rules for introducing mismatched bases are as follows: a “strong” mismatch at the 3’ end of the specific editing primer requires a “weak” mismatch at the second position, and vice versa; while a “medium” mismatch at the 3’ end should be paired with a “medium” second mismatch (22, 24) (Table S2). Such intentional mismatches can increase the quantification cycle (Ct) by approximately 3–9 cycles, thereby enabling effective discrimination between edited and unedited templates while maintaining measurable amplification for quantitative analysis. The reaction system and protocol are performed on the Bio-Rad CFX Connect™ Real-Time System (Bio-Rad, USA). The reaction mixture is 20 µL, containing Hieff® qPCR SYBR® Green Master Mix (YEASEN, 11201ES05, Shanghai, China), 100 ng cDNA, and 10 µM of primers. The PCR conditions (three-step protocol) are as follows: 95°C for 5 minutes for pre-denaturation, followed by 41 cycles of 95°C for 10 seconds, 62°C for 20 seconds, and 72°C for 20 seconds.

For the RES-qPCR results, after the reaction is complete, the amplification products are analyzed by 1.5% agarose gel electrophoresis and Sanger sequencing. The primers that meet the expected criteria are selected for subsequent data quantification. The standards are as follows: (1) When cDNA is used as the template, both the unedited- and edited-stie primers should amplify normally, producing a single band, while no amplification product should be observed when gDNA is used as the template; (2) Sanger sequencing results should confirm the presence of both the edited and unedited transcript forms, along with the introduced mismatch bases.

The relative expression levels of RNA-editing transcripts are calculated using the 2^-△Ct^ method, where the relative expression of RNA-editing transcripts is normalized to the gene expression level of the RES in the sample, while other data are normalized to the expression level of the nuclear elongation factor 1-alpha (*EF1*α) gene.

### Gene expression quantification

Transcript abundance was also quantified as TPM (transcripts per million) using TPMCalculator (v0.0.3) (25). BAM files generated by RMTA and the corresponding GTF annotation files were provided to TPMCalculator (-b for BAM, -g for annotation).

### Analysis of RNA editing enzymes

Protein annotation files of 21 arthropod species were retrieved from the NCBI public database (Table S3). The Pfam-A database (downloaded on 2025-08-03) was used to identify conserved domains. In total, 35 relevant domains were iok selected, including members of the deaminase superfamily clan CL0109 (33 families), the double-stranded RNA-binding domain (PF00035), and Z-DNA/RNA-binding domains (PF02295 and PF03178) (Dataset S4). Domain annotation was performed with hmmscan in HMMER (version 3.2) (parameters: --cut_ga, domain E-value ≤ 1e-5, coverage ≥ 0.5) (26), retaining only the highest-scoring annotation in cases of overlapping hits. Candidate sequences were further validated with InterProScan to refine domain boundaries and eliminate false positives (27). For genes with multiple isoforms, only the longest transcript was retained, and truncated sequences shorter than 100 amino acids were discarded. Multiple sequence alignments of candidate proteins were generated with MUSCLE (version 5.3) (28), and alignments were trimmed with trimAl (version 1.5.rev0) (-automated1) (29) to remove highly gapped columns and low-quality sequences. Phylogenetic trees were constructed with IQ-TREE (version 2.4.0) using ModelFinder (-m MFP) and branch supports were assessed with 1,000 ultrafast bootstrap replicates (-bb 1000) and 1,000 SH-aLRT replicates (-alrt 1000) (69). Finally, the resulting trees were visualized and refined with iTOL (https://itol.embl.de/) (30).

### Double-stranded RNA synthesis

Double-stranded RNA (dsRNA) of *ADAR1* and *ADAR2* was synthesized in vitro using the T7 High Yield RNA Transcription Kit (Vazyme, DD4201, Nanjing, China) according to the manufacturer’s instructions. Gene-specific primers containing the T7 polymerase promoter sequence were designed using the DSIR website (http://biodev.extra.cea.fr/DSIR/DSIR.html). All synthesized dsRNAs were dissolved in nuclease-free water and quantified by a K5600C micro-volume spectrophotometer (KAIAO, Beijing, China)

### RNAi experiments

The feeding treatment was carried out according to the method described in (63), with minor modifications. Double-layered stretched parafilm (Parafilm, PM996, USA) was used to cover acrylic rings (25 mm in diameter × 15 mm in height), constructing the feeding apparatus. Each device was filled with 50 μL of artificial feed mixed with dsRNA, which included a final concentration of 30% sucrose solution, 0.02% neutral red dye, and 1 μg/μL of dsRNA.

To assess the impact of ADARs on *M. persicae* host colonization, 20 synchronized third-instar GPA reared on Bn were fed with *dsADARs* (*dsADAR1* and *dsADAR2*) for 48 hours. The surviving aphids were then transferred to Bn plants for a 10-day reproductive capacity assessment. *dsLta* (Lymphotoxin A gene of *Mus musculus*) was used as a control (33). The interference efficiency was evaluated via qPCR. Each biological replicate was independently tested three times (n = 3), and for each biological replicate, two technical replicates were performed.

### Host plant change experiments

Approximately 150 adult aphids from GPA-Bn and GPA-Zm were collected and introduced onto 4-week-old host plants. After 24 hours, adults were removed, while the newly hatched nymphs remained on the plants for three days. Thirty synchronized 3-day-old aphids were then transferred simultaneously to both their original and novel host plants. This transfer procedure was repeated at least three times, with each transfer serving as an independent biological replicate. To prevent escape, aphids were confined to the plant leaves within foam clip-cages (2 cm height × 3.5 cm diameter), securely sealed at the base of the petiole. Two days after the transfer, live nymphs from both original and novel plants were collected for RNA extraction. Each biological replicate was subjected to two technical replicates of quantitative reverse transcription PCR (qRT-PCR) analysis.

To investigate the effects of gene silencing of *ADARs* on RNA editing and rapid host adaptation of *M. persicae*, GPA-Bn 3-day-old nymphs were fed *dsADAR1* and *dsADAR2* (1 μg/μL) for 48 hours. Control groups included nymphs fed with *dsLta*. After the feeding period, the same transfer procedure was performed, and subsequent experiments included: (a) quantification of candidate RNA editing site (RES) levels and related gene expression; (b) measurement of fecundity and survival rate over 10 days. All experiments were performed independently three times, and for each biological replicate, two technical replicates were included for quantitative measurements.

### Statistical analysis

All analyses were performed in R. Two-group comparisons (e.g., dsRNA-treated vs. control) for RNA editing levels, transcript ratios, gene expression, and fecundity were conducted using Wilcoxon rank-sum tests. Differences in ADAR TPM across hosts were analyzed by one-way ANOVA, with normality and homogeneity of variances assessed using the Shapiro–Wilk and Levene’s tests, respectively; post-hoc pairwise comparisons were performed using Tukey’s HSD or Games–Howell as appropriate. Survival curves were compared using the log-rank test. Host-dependent RNA editing per site was assessed by linear regression (lm(value ∼ hosts)), with ANOVA F-tests determining significance (*P* < 0.05 and fold-change ≥ 2). The distribution of A-to-I RNA editing sites across genomic regions and the proportions of synonymous and nonsynonymous sites within CDS regions were evaluated using Pearson’s chi-square tests, with post-hoc pairwise comparisons where appropriate

---

**Fig. S1.**
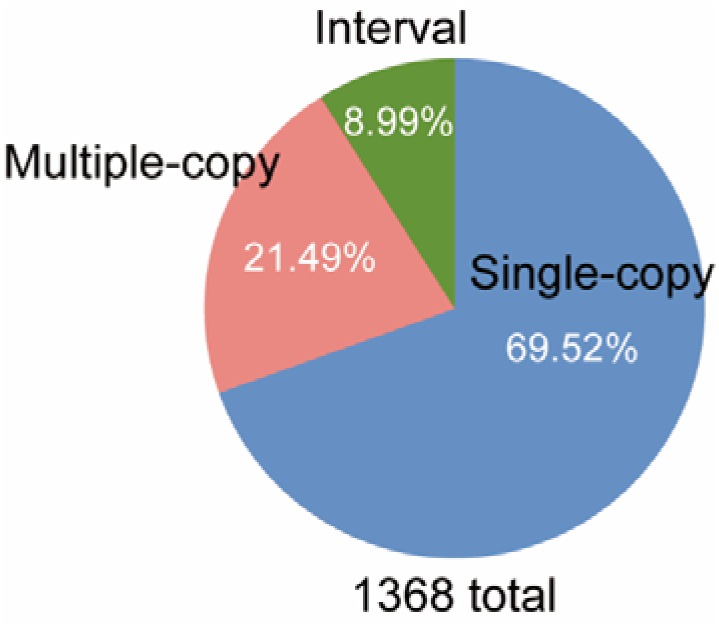
Proportional distribution of candidate RES across single-copy, multi-copy, and intergenic regions. The pie chart shows the proportion of 1,368 sites with percentages indicated. Subsequent analyses focused only on single-copy genes to avoid potential artifacts from unassembled duplicates or tandem repeats.

**Fig. S2.**
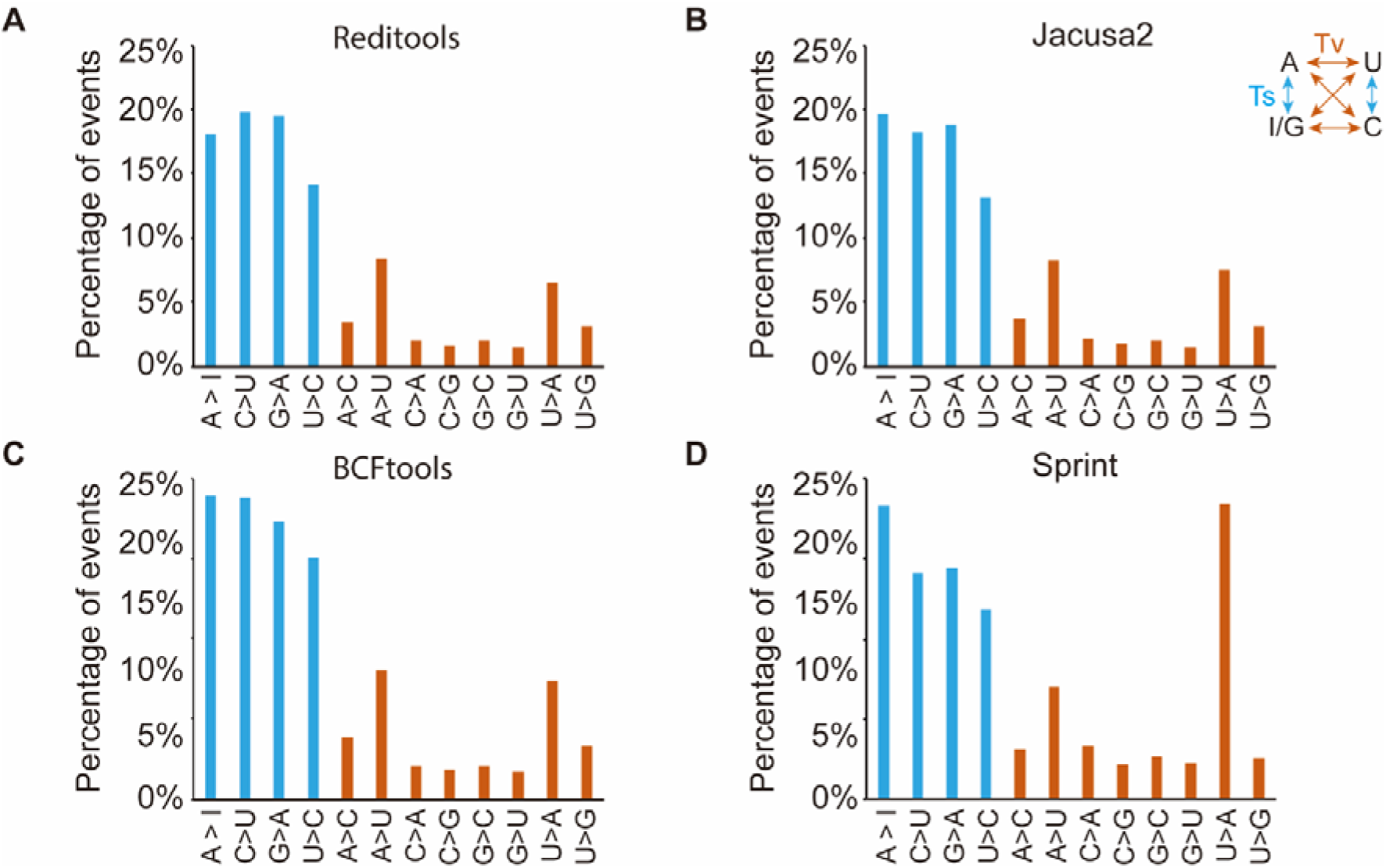
Distribution of the 12 possible nucleotide substitution types among all RES identified by four bioinformatic software tools. The 12 substitution types were counted based on RES detected by the four tools following the workflow in Fig. 1D, showing that the results are largely consistent with those in Fig. 1F.

**Fig. S3.**
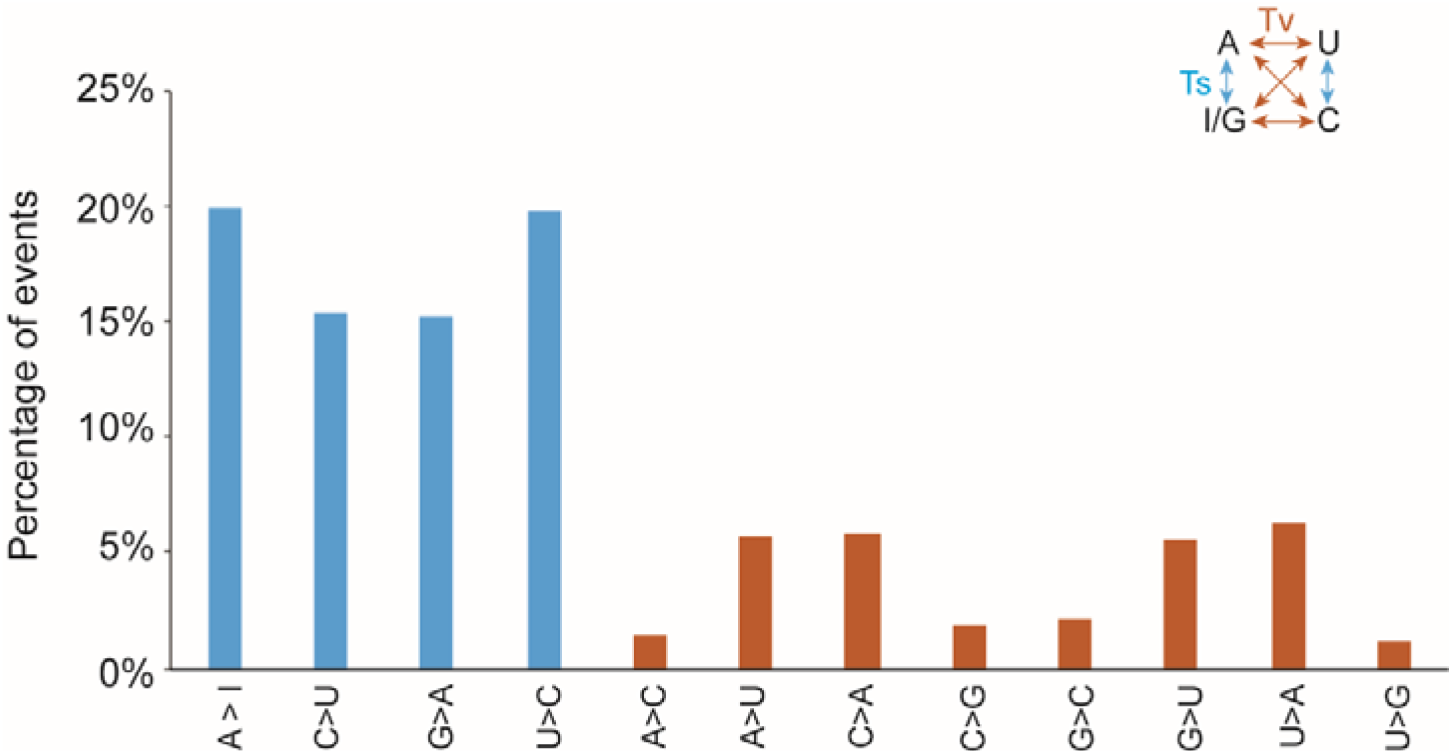
Proportion of 12 types of RES. RNA-seq data (accession: ERR1661483) (1) from *M. persicae* Clone O reared on *B. rapa* was used for the analysis. RES were identified using Reditools.

**Fig. S4.**
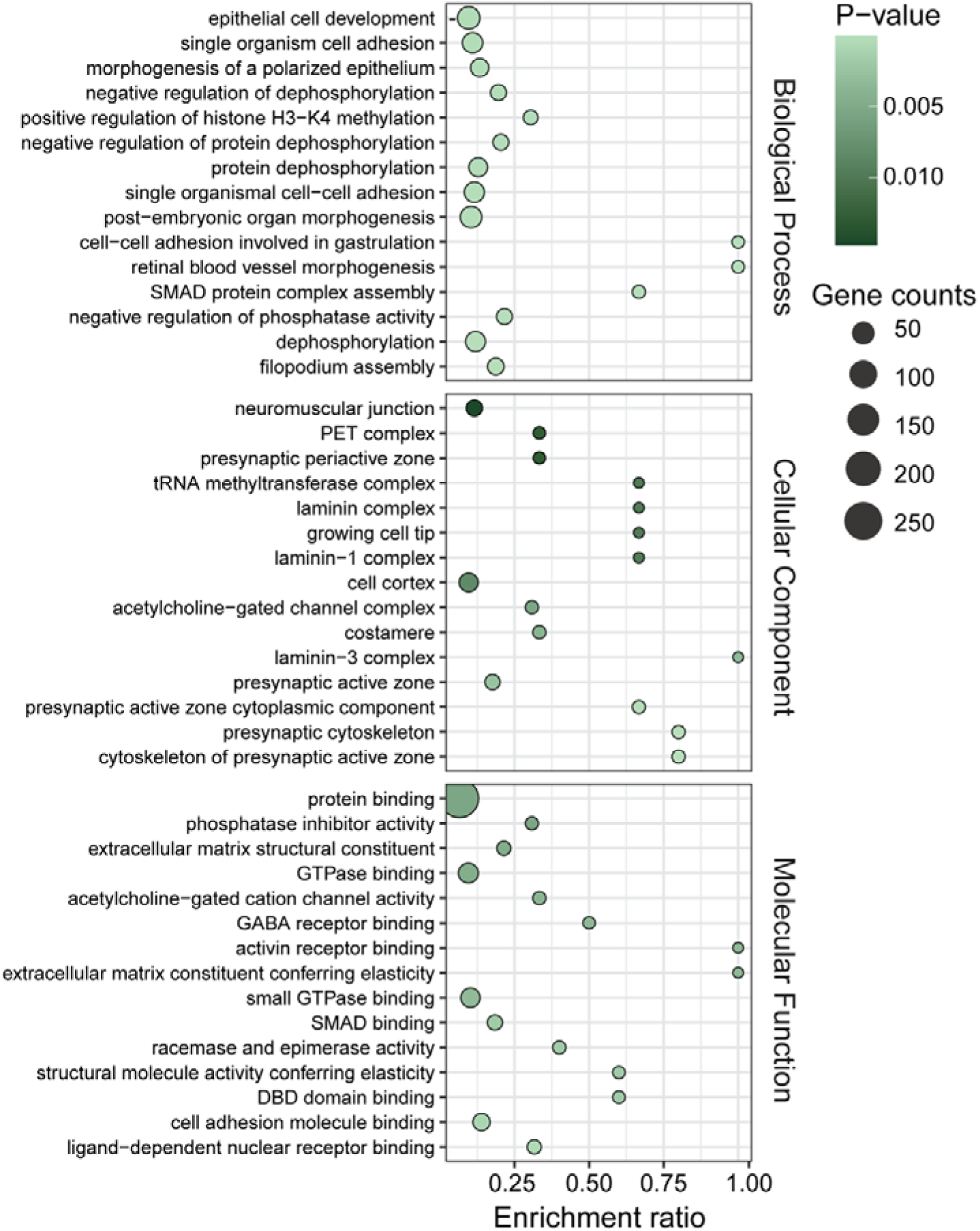
GO term enrichment analysis of genes within RESs located coding region.

**Fig. S5.**
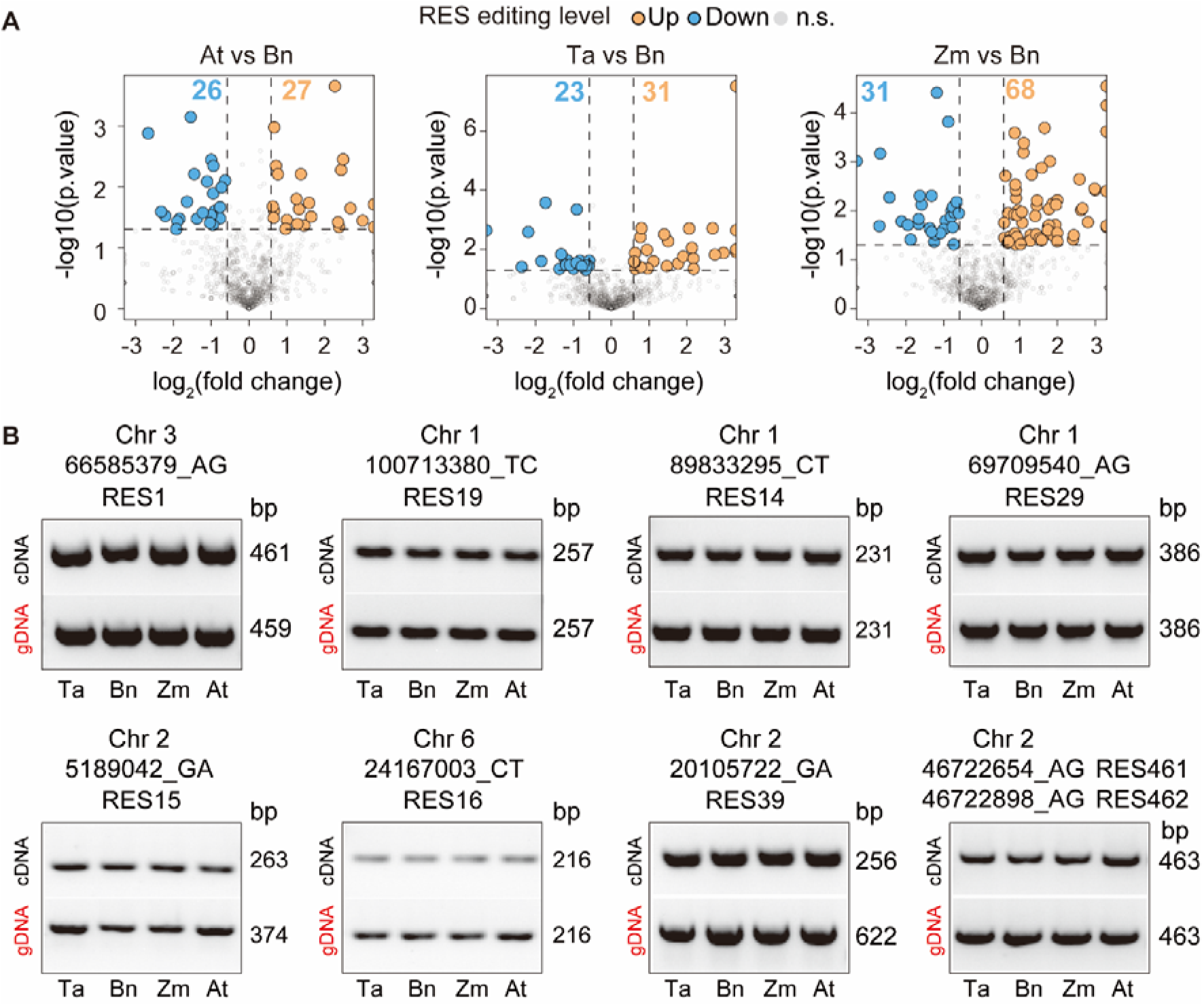
RES are differentially edited in *M. persicae* on different plant hosts. (A) Volcano plot showing differential RNA editing between aphids reared on various host plants. Editing levels in aphids on Bn were used as the control. DE RES were defined as those with a ≥2-fold change in editing level and *P* < 0.05 (ANOVA F-test). (B) Gel electrophoresis image of PCR amplicons containing selected RES. Forward and reverse primers were designed to flank the target RES. PCR was performed using both cDNA and genomic DNA (gDNA) templates, followed by Sanger sequencing.

**Fig. S6.**
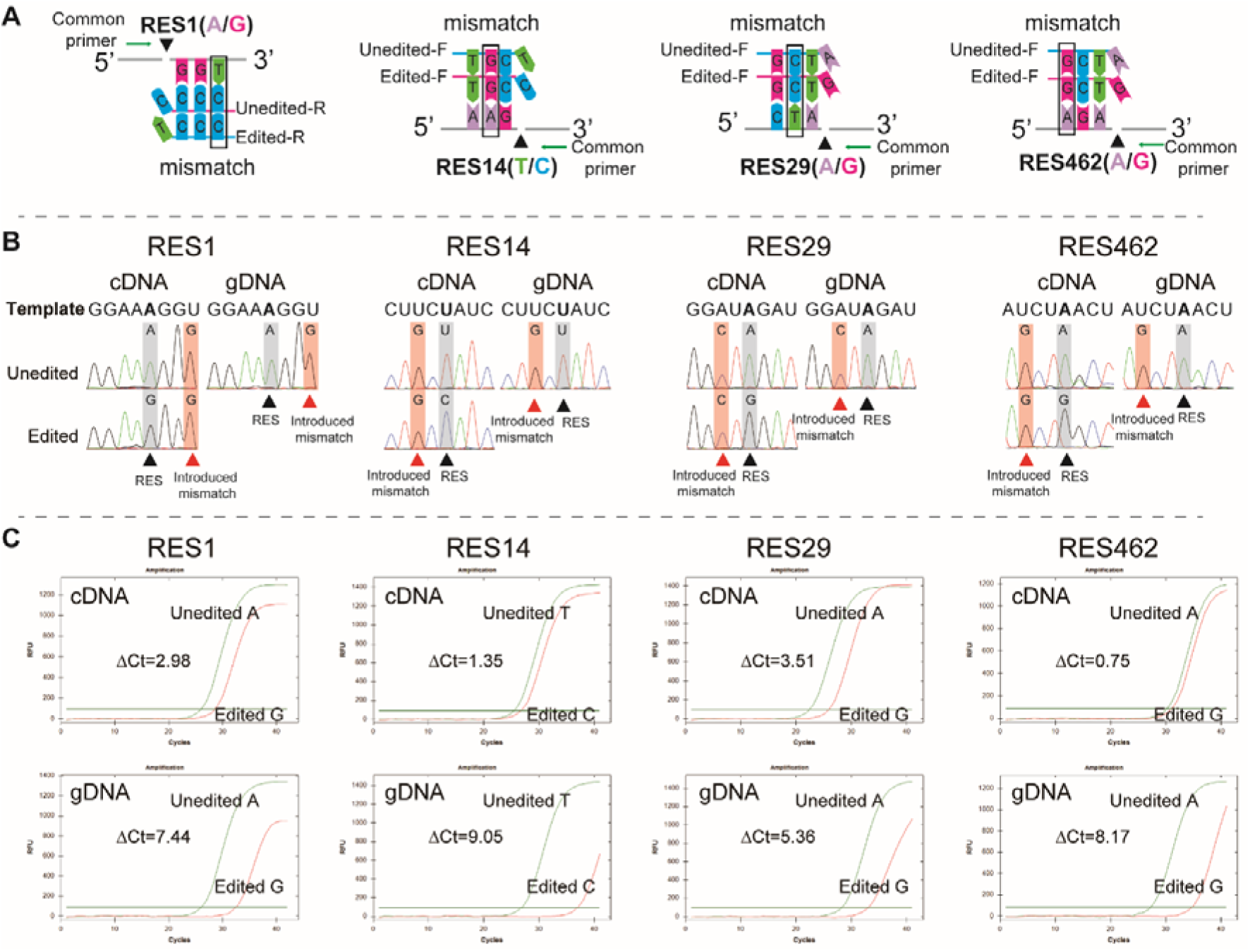
Validation of RES-qPCR for distinguishing edited and unedited transcripts. (A) Schematic illustration of RES-qPCR primer design. The forward primers contain the edited or unedited nucleotide at the 3′ end and include an additional mismatch 1–2 bases upstream (black boxes), paired with a common reverse primer. (B) Representative Sanger sequencing chromatograms of PCR products amplified from cDNA and gDNA templates for four selected RES (RES1, RES14, RES29, and RES462). Arrows indicate RES (black) and the introduced mismatch nucleotide (red). (C) Amplification curves of edited and unedited primers using cDNA and gDNA templates. Green curves correspond to amplification with unedited primers, and red curves correspond to amplification with edited primers. ΔCt values indicate differences in amplification efficiency between the two primer sets. In cDNA templates containing edited transcripts, the assay clearly distinguishes between edited and unedited transcripts. In contrast, amplification from gDNA templates is strongly suppressed due to 3′-end mismatches and the presence of artificially introduced sequence alterations.

**Fig. S7.**
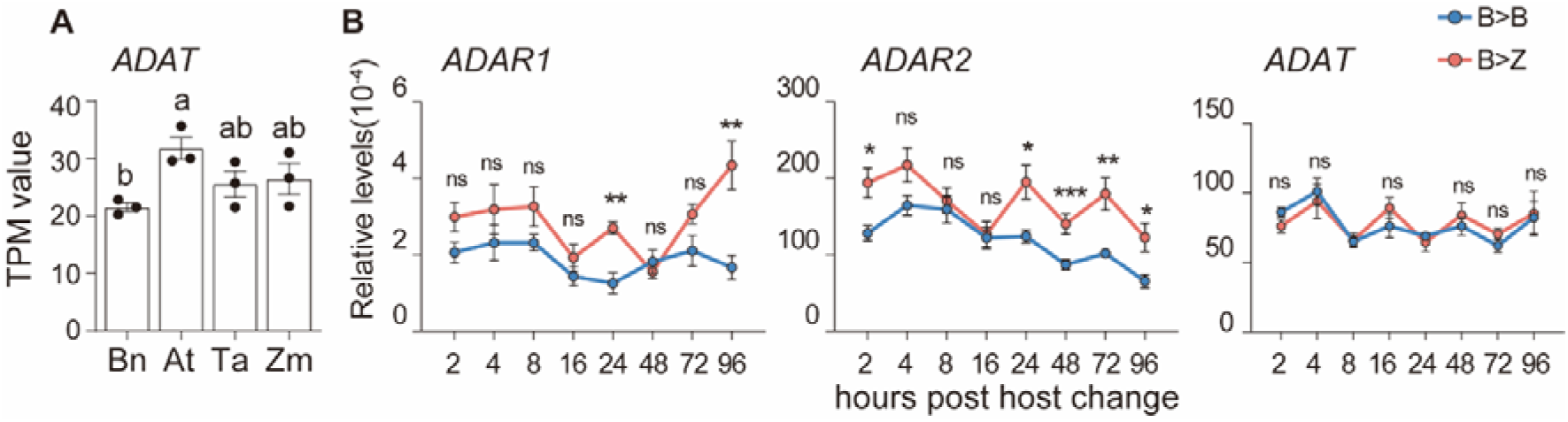
Dynamic gene expression and RNA editing in aphids after host transfer. (A) TPM of *ADAT* in *M. persicae* reared on different host plants. Different lowercase letters indicate significant differences (one-way ANOVA with Tukey’s test). (B) Expression levels of *ADAR1*, *ADAR2* and *ADAT* at different time points after transfer from Bn to Zm. Data are presented as mean ± SEM (n = 6; 15 aphids per biological replicate). Symbols indicate statistical significance: *P* < 0.05, *P* < 0.01, *P* < 0.001; ns = not significant. Blue lines: B > B; red lines: B >Z.

**Fig. S8.**
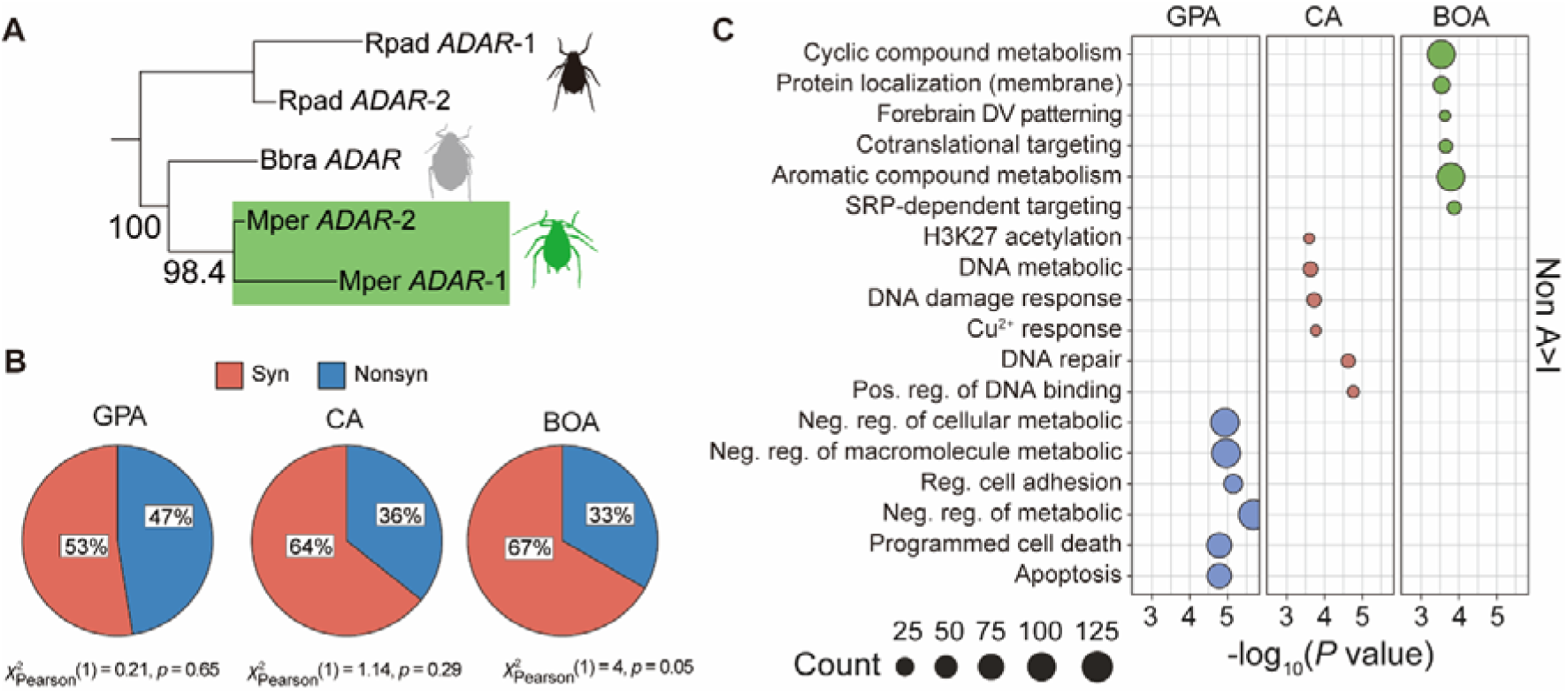
Dynamic gene expression and RNA editing in aphids after host transfer. (A) Phylogeny of ADAR homologs from *Myzus persicae* (Mper), *Rhopalosiphum padi* (Rpad), and *Brevicoryne brassicae* (Bbra) (B) Proportions of synonymous (Syn) and nonsynonymous (NonSyn) A-to-I editing sites within CDS regions in the three aphid species. Pearson’s chi-square test results were used to evaluate differences in proportions among the aphid species. (C) Gene Ontology (GO) enrichment analysis of genes harboring non-A-to-I RES. GO terms were detailed in Table S4. Enriched biological processes are plotted based on gene count (dot size) and significance (−log_10_ (*P value*))

**Table S1.**
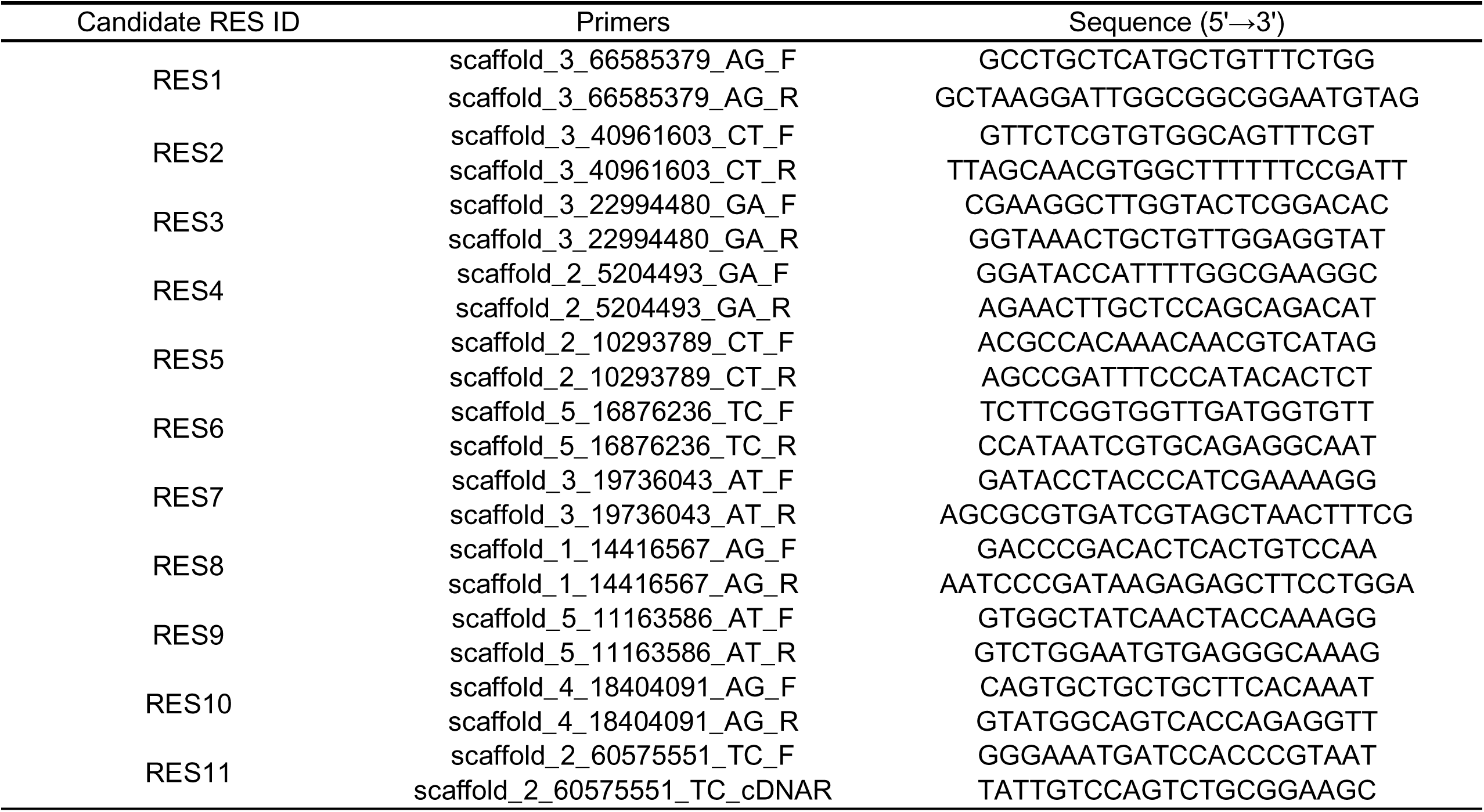
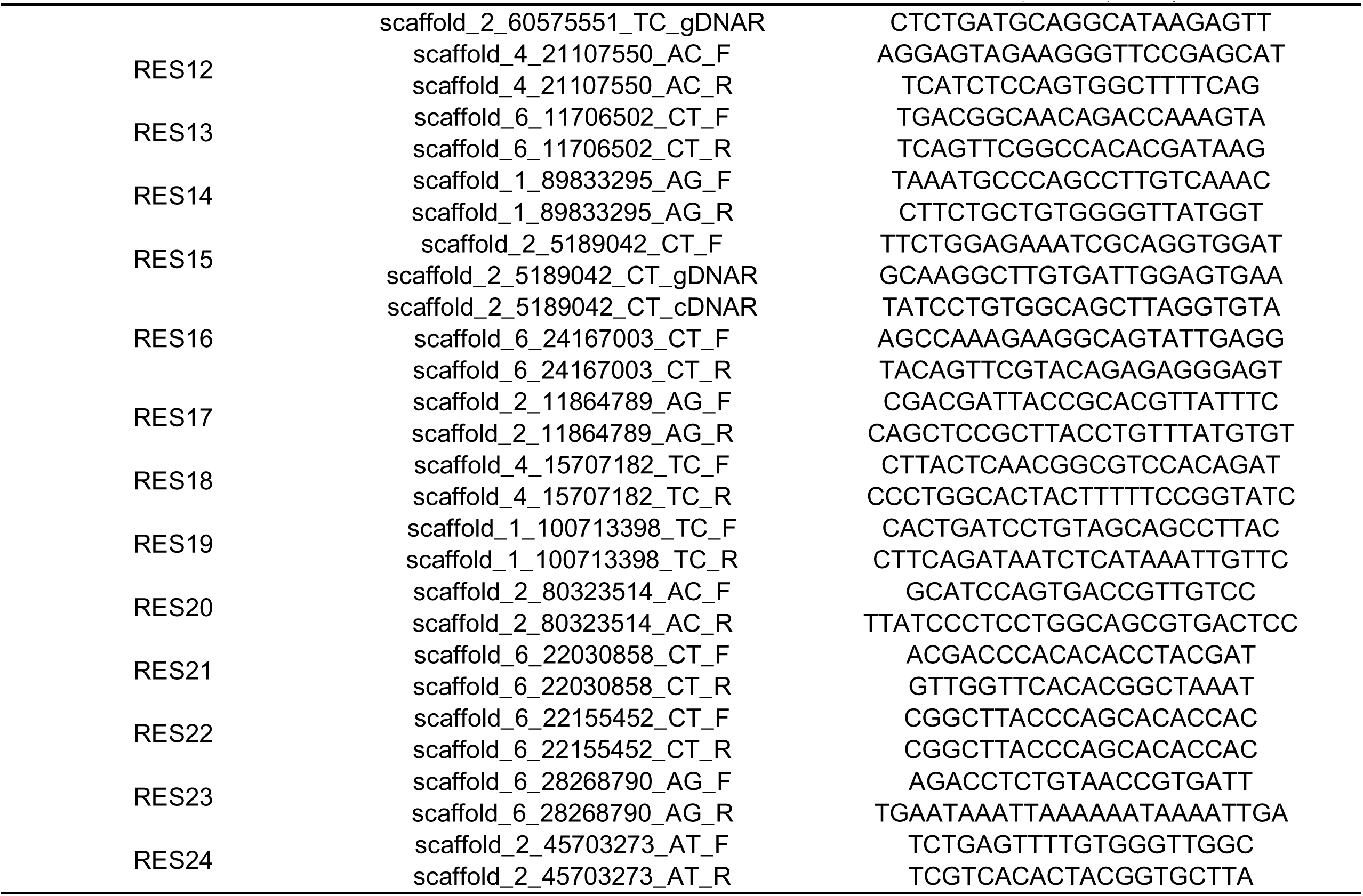
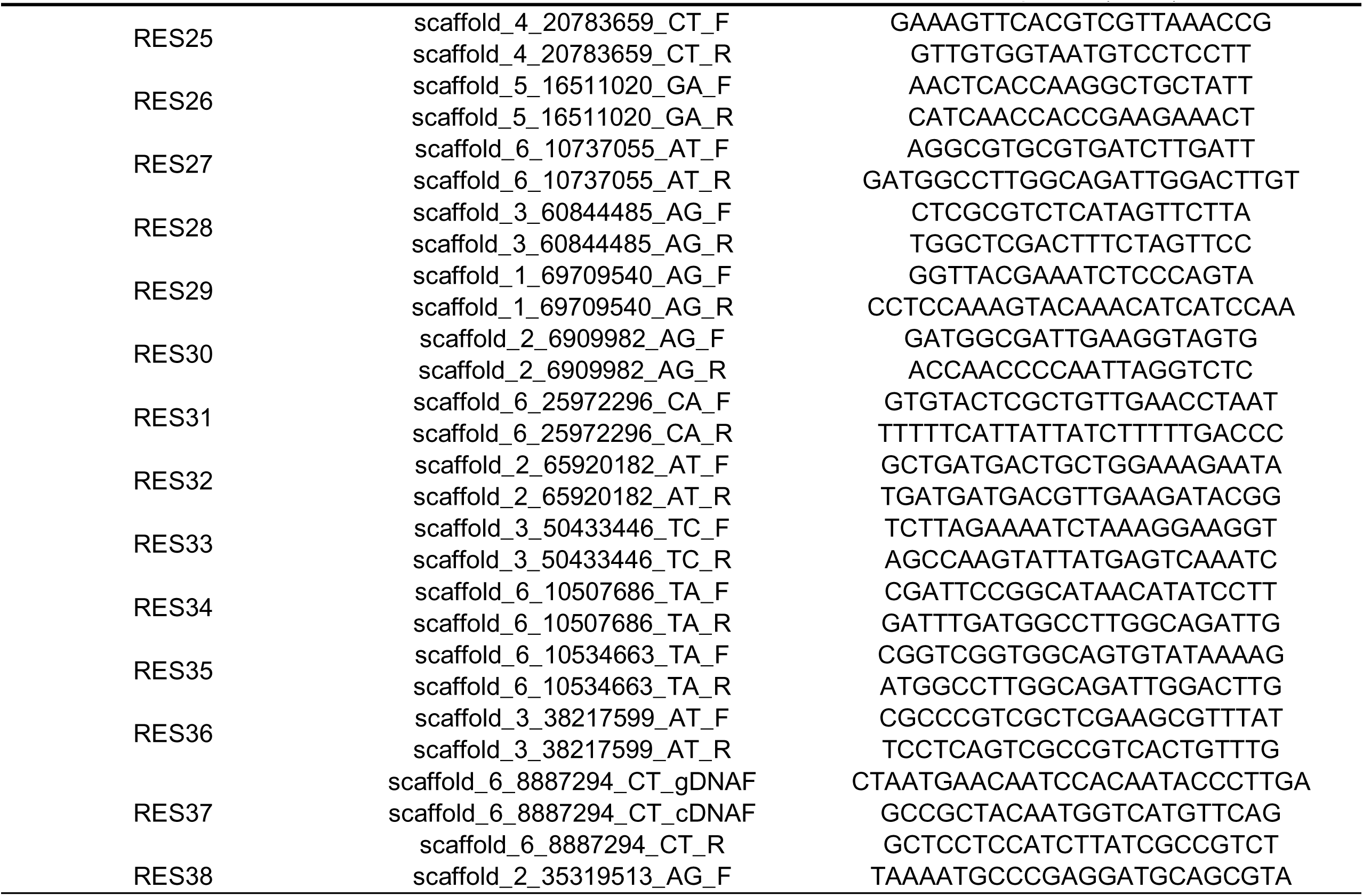
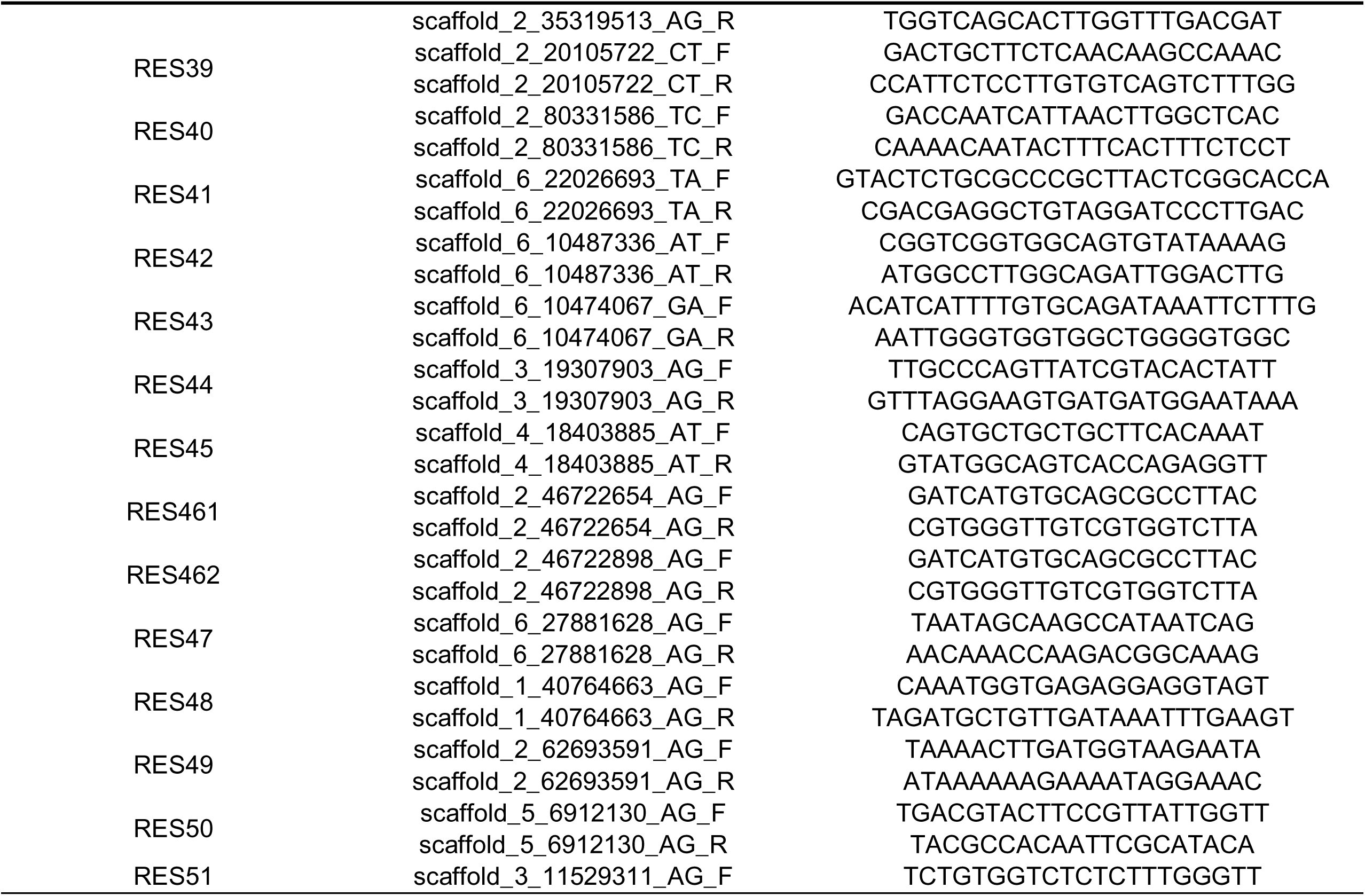
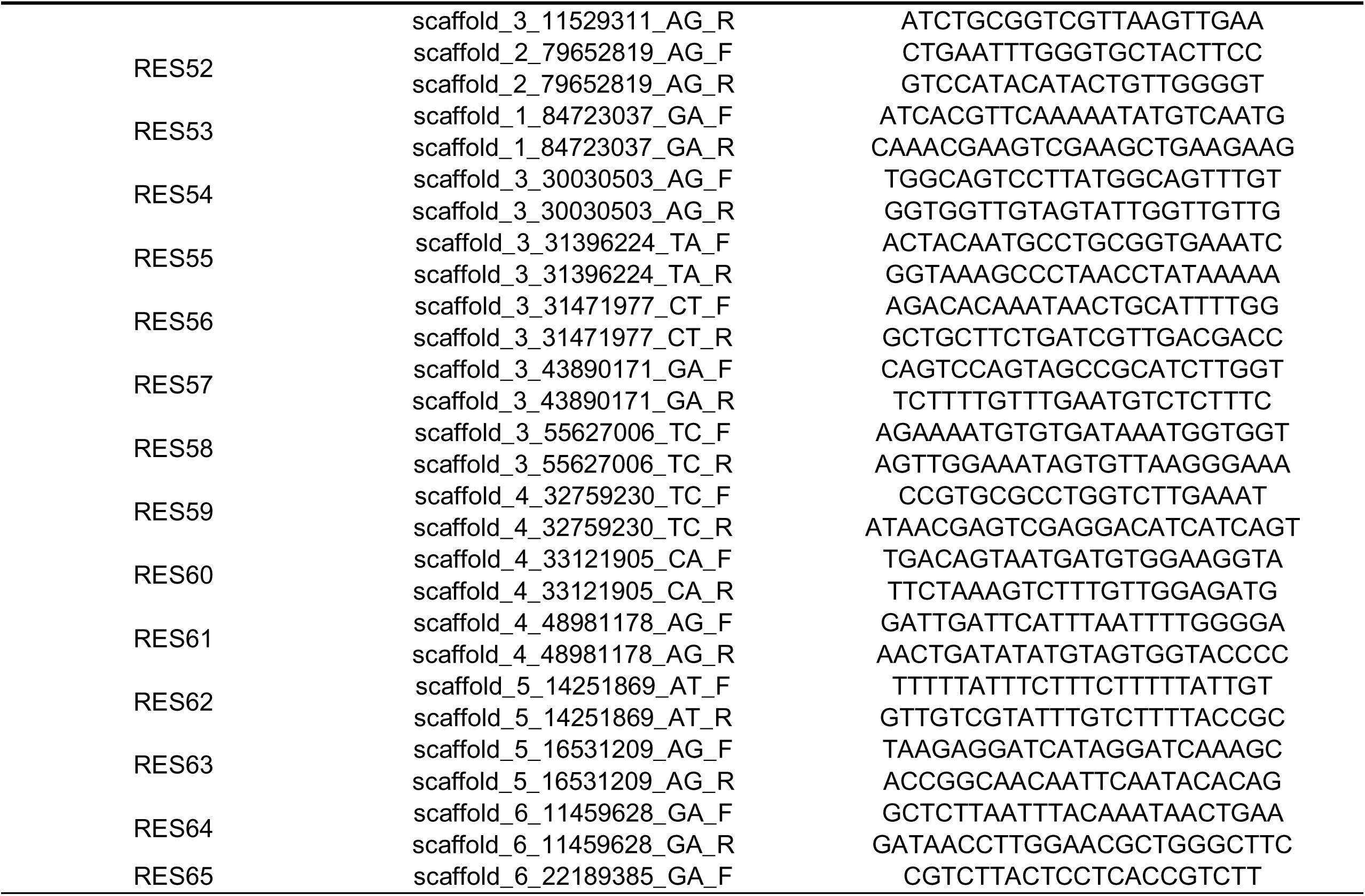
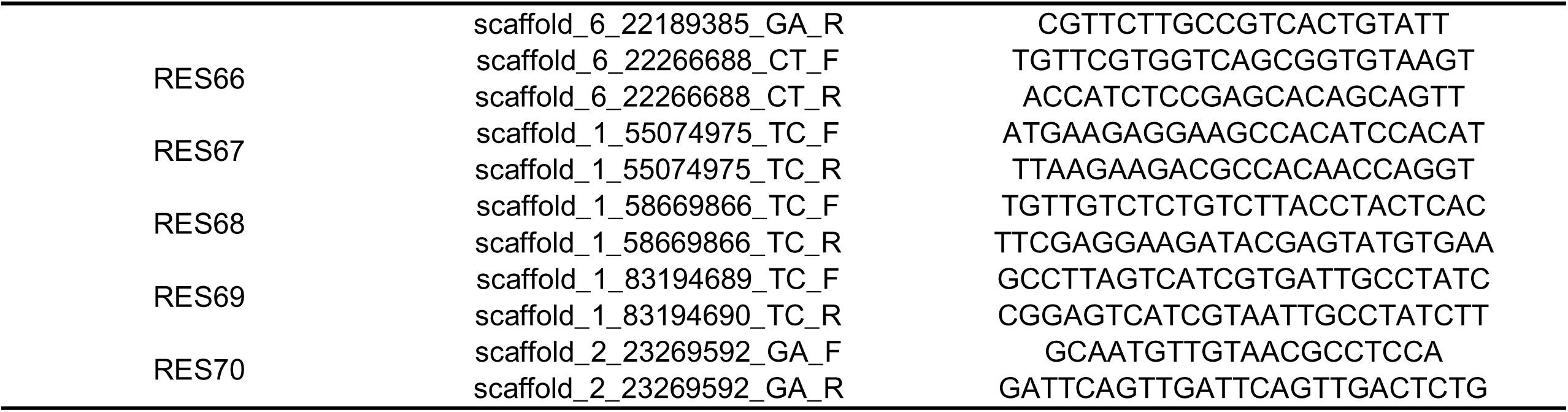
Primers used for validation of candidate RNA editing sites (RESs). Primers were used to amplify regions containing candidate RESs using cDNA and genomic DNA (as a control) as templates, in der to validate the presence of RNA editing.

**Table S2.**
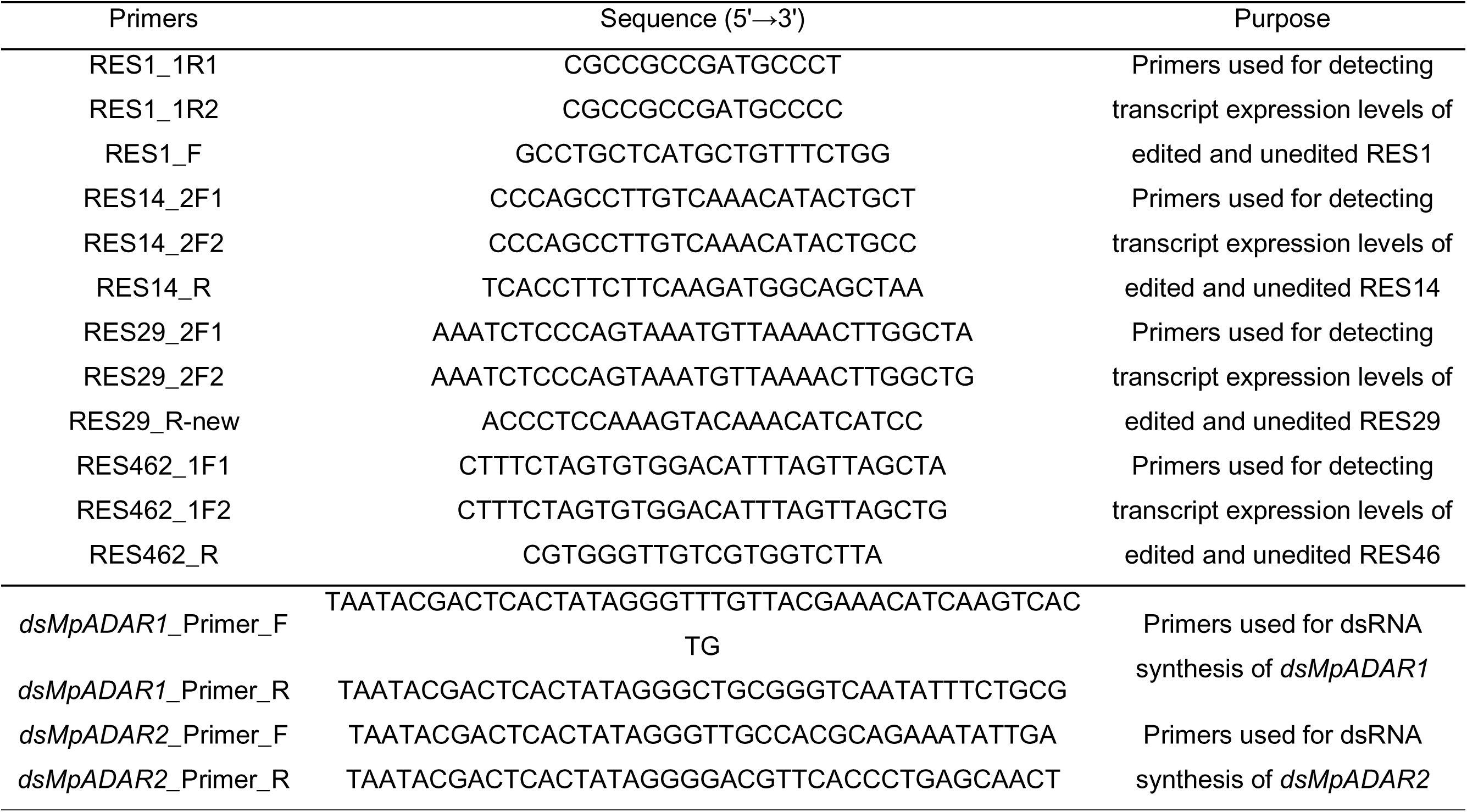
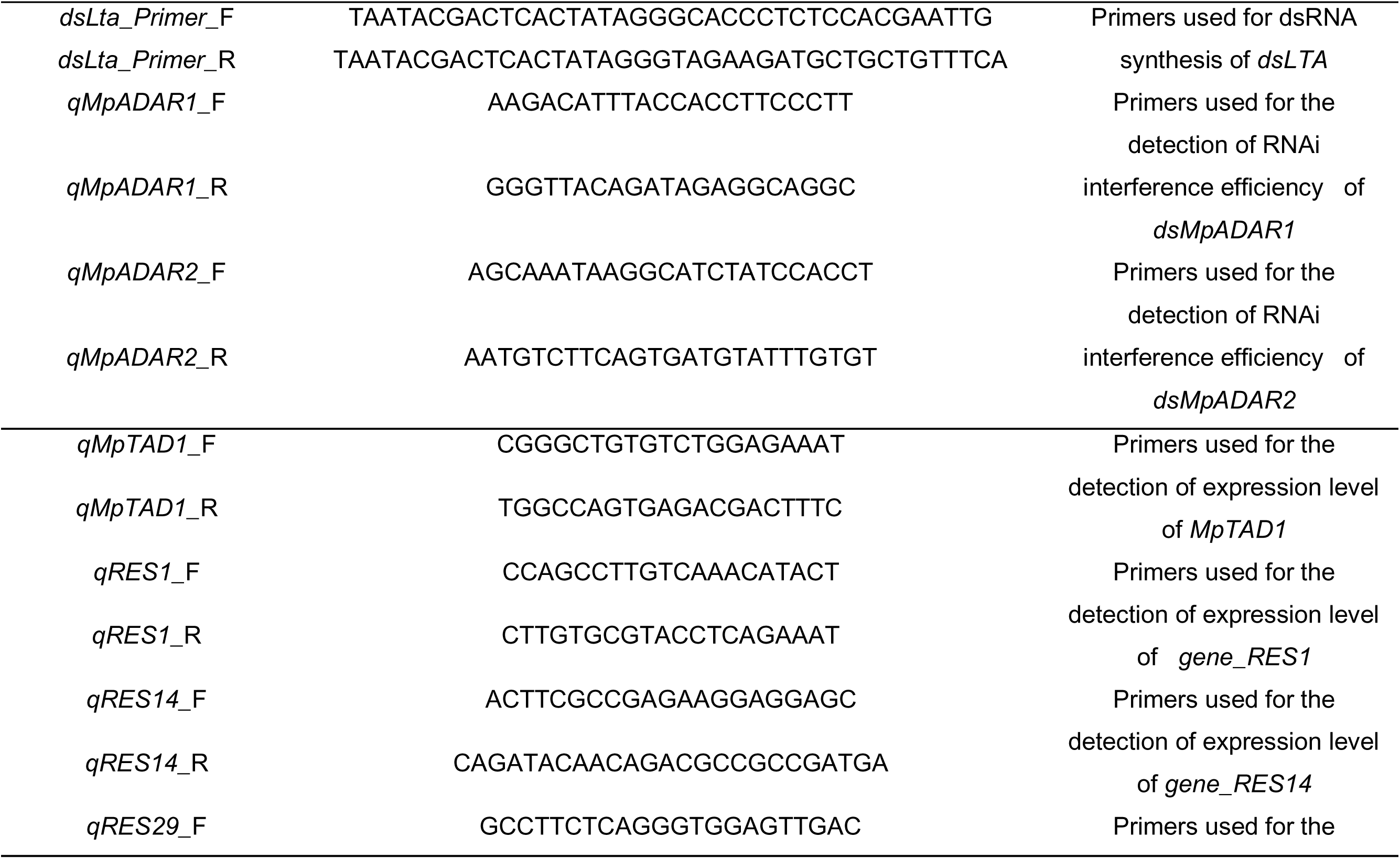
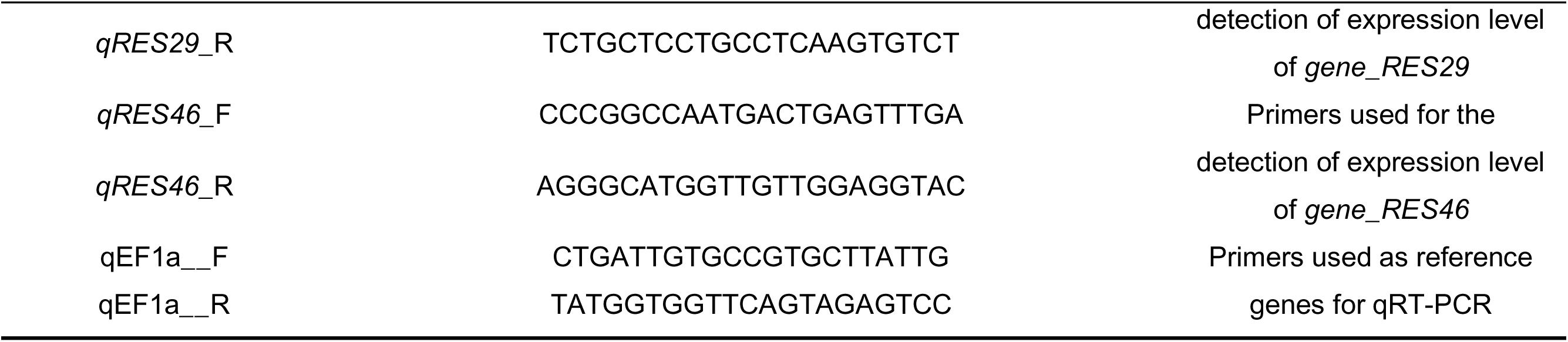
List of primers used in this study and their experimental purposes. . Primers were designed for validation of candidate RNA editing sites (RESs), quantification of edited and unedited transcript isoforms, dsRNA synthesis for RNA interference, and qRT-PCR assays of gene expression or RNAi efficiency.

**Table S3.** Genome data source of 22 species used in Phylogenetic analysis of ADARs and ADAT. .

| Species | Order | family | Genus | Abbreviation in tree | Source |
| --- | --- | --- | --- | --- | --- |
| *Tetranychus urticae* | Trombidiformes | Tetranychidae | *Tetranychus* | Turt | GRBI |
| *Atta cephalotes* | Hymenoptera | Formicidae | *Atta* | Acep | GCF_000143395.1 |
| *Tribolium castaneum* | Coleoptera | Tenebrionidae | *Tribolium* | Tcas | GCF_031307605.1 |
| *Daktulosphaira vitifoliae* | Hemiptera | Phylloxeridae | *Daktulosphaira* | Dvit | Aphid Base |
| *Myzus persicae* | Hemiptera | Aphididae | *Myzus* | Mper | (3) |
| *Acyrtosiphon pisum* | Hemiptera | Aphididae | *Acyrtosiphon* | Apis | (34) |
| *Rhopalosiphum padi* | Hemiptera | Aphididae | *Rhopalosiphum* | Rpad | (3) |
| *Rhopalosiphum maidis* | Hemiptera | Aphididae | *Rhopalosiphum* | Rmai | (35) |
| *Aphis gossypii* | Hemiptera | Aphididae | *Aphis* | Agos | (3) |
| *Aphis glycines* | Hemiptera | Aphididae | *Aphis* | Agly | (36) |
| *Melanaphis sacchari* | Hemiptera | Aphididae | *Aphis* | Msac | GCF_002803265.2 |
| *Bemisia tabaci* | Hemiptera | Aleyrodidae | *Bemisia* | Whitefly | GCF_001854935.1 |
| *Rhodnius prolixus* | Hemiptera | Reduviidae | *Rhodnius* | Rpro | (37) |
| *Oncopeltus fasciatus* | Hemiptera | Lygaeidae | *Oncopeltus* | Ofas | (38) |
| *Frankliniella occidentalis* | Thysanoptera | Thripidae | *Frankliniella* | Focc | (39) |
| *Daphnia pulex* | Cladocera | Daphniidae | *Daphnia* | Dpul | GCA_000187875.1 |
| *Leptinotarsa decemlineata* | Coleoptera | Chrysomelidae | *Leptinotarsa* | Ldec | NCBI genome |
| *Myzus cerasi* | Hemiptera | Aphididae | *Myzus* | Mcer | Aphid Base |
| *Schlechtendalia chinensis* | Hemiptera | Aphididae | *Schlechtendalia* | Schi | GCA_019022885.1 |
| *Sipha flava* | Hemiptera | Aphididae | *Sipha* | Sfla | GCA_003268045.1 |
| *Sitobion miscanthi* | Hemiptera | Aphididae | *Sitobion* | Smis | GCA_008086715.1 |
| *Plutella xylostella* | Lepidoptera | Plutellidae | *Plutella* | Pxyl | GCF_000330985.1 |

**Table S4.** Detailed GO enrichment analysis of genes harboring RES identified in the three aphid species under host-shift conditions.

| Description | Count | GeneRatio | Pvalue | Species | type |
| --- | --- | --- | --- | --- | --- |
| cell-type specific apoptotic process | 18 | 0.058 | 6.22E-06 | GPA | A>I |
| apoptotic process | 35 | 0.035 | 3.94E-05 | GPA | A>I |
| programmed cell death | 36 | 0.033 | 7.02E-05 | GPA | A>I |
| regulation of epithelial cell differentiation | 8 | 0.096 | 8.92E-05 | GPA | A>I |
| glial cell fate determination | 3 | 0.5 | 9.87E-05 | GPA | A>I |
| neuron apoptotic process | 11 | 0.061 | 0.000269 | GPA | A>I |
| white fat cell differentiation | 2 | 0.118 | 0.000726 | CA | A>I |
| regulation of chondrocyte differentiation | 2 | 0.083 | 0.001459 | CA | A>I |
| regulation of multicellular organismal development | 10 | 0.006 | 0.00165 | CA | A>I |
| positive regulation of histone modification | 3 | 0.028 | 0.002101 | CA | A>I |
| regulation of developmental process | 11 | 0.006 | 0.002162 | CA | A>I |
| maintenance of imaginal histoblast diploidy | 1 | 1 | 0.002396 | CA | A>I |
| negative regulation of chromatin binding | 3 | 0.5 | 1.04E-05 | BOA | A>I |
| proton-transporting V-type ATPase complex assembly | 2 | 0.5 | 0.000396 | BOA | A>I |
| vacuolar proton-transporting V-type ATPase complex assembly | 2 | 0.5 | 0.000396 | BOA | A>I |
| protein targeting | 11 | 0.026 | 0.000564 | BOA | A>I |
| protein targeting to membrane | 6 | 0.047 | 0.000632 | BOA | A>I |
| female germ-line sex determination | 2 | 0.4 | 0.000656 | BOA | A>I |
| negative regulation of metabolic process | 131 | 0.065 | 2.07E-06 | GPA | Non A>I |
| regulation of cell adhesion | 31 | 0.107 | 7.11E-06 | GPA | Non A>I |
| negative regulation of macromolecule metabolic process | 118 | 0.065 | 1.10E-05 | GPA | Non A>I |
| negative regulation of cellular metabolic process | 114 | 0.065 | 1.18E-05 | GPA | Non A>I |
| programmed cell death | 78 | 0.072 | 1.66E-05 | GPA | Non A>I |
| apoptotic process | 74 | 0.073 | 1.66E-05 | GPA | Non A>I |
| positive regulation of DNA binding | 4 | 0.105 | 1.73E-05 | CA | Non A>I |
| DNA repair | 9 | 0.023 | 2.36E-05 | CA | Non A>I |
| cellular response to copper ion | 2 | 0.4 | 0.00017 | CA | Non A>I |
| cellular response to DNA damage stimulus | 10 | 0.016 | 0.000189 | CA | Non A>I |
| DNA metabolic process | 11 | 0.014 | 0.000234 | CA | Non A>I |
| histone H3-K27 acetylation | 2 | 0.333 | 0.000254 | CA | Non A>I |
| SRP-dependent cotranslational protein targeting to membrane | 8 | 0.121 | 0.000133 | BOA | Non A>I |
| cellular aromatic compound metabolic process | 106 | 0.03 | 0.000164 | BOA | Non A>I |
| cotranslational protein targeting to membrane | 8 | 0.113 | 0.000224 | BOA | Non A>I |
| forebrain dorsal/ventral pattern formation | 3 | 0.5 | 0.000235 | BOA | Non A>I |
| protein localization to membrane | 21 | 0.054 | 0.000287 | BOA | Non A>I |
| organic cyclic compound metabolic process | 107 | 0.03 | 0.000294 | BOA | Non A>I |

**Table S5.** Detailed annotation of candidate RNA-editing sites (RESs).

| Candidate<br>RES ID | Site | Strand | gene_id | Genomic<br>feature | Annotation |
| --- | --- | --- | --- | --- | --- |
| RES1 | scaffold_3_66585379_AG | + | MYZPE13164_O_Elv2.1_0259980 | CDS | PREDICTED: *Myzus persicae* potassium voltage-gated channel protein Shab (LOC111031521) |
| RES29 | scaffold_1_69709540_AG | + | MYZPE13164_O_Elv2.1_0085790 | 3'-UTR | Zinc finger BED domain-containing protein 4-like |
| RES462 | scaffold_2_46722898_AG | + | MYZPE13164_O_Elv2.1_0171060 | 3'-UTR | PREDICTED *Myzus persicae* origin recognition complex subunit 3 (LOC111040050) |
| RES14 | scaffold_1_89833295_AG | - | MYZPE13164_O_Elv2.1_0099850 |  | PREDICTED: *Myzus persicae* uncharacterized LOC111038309, ncRNA |

